# A diencephalic pathway for movement initiation and rescue of Parkinsonian symptoms

**DOI:** 10.1101/395277

**Authors:** Glenn D.R. Watson, Ryan N. Hughes, Elijah A. Petter, Henry H. Yin

## Abstract

The parafascicular nucleus (Pf) of the thalamus projects to the subthalamic nucleus (STN), a major target for deep brain stimulation (DBS) in Parkinson’s disease (PD), but the function of this projection remains unknown. Here, we used optogenetics, 3D motion capture, *in vivo* electrophysiology and 1-photon calcium imaging, unsupervised behavioral classification, and viral-based neuroanatomical tracing to examine the contribution of Pf efferents to movement generation in mice. We discovered that Pf neurons are highly correlated with movement velocity and excitation of Pf neurons generates turning and orienting movements. Movement initiation was not due to Pf projections to the striatum, but rather its projections to the STN. Optogenetic excitation of the Pf-STN pathway restores movement in a common mouse model of PD with complete akinesia. Collectively, our results reveal a thalamo-subthalamic pathway regulating movement initiation, and demonstrate a circuit mechanism that could potentially explain the clinical efficacy of DBS for relief of PD motor symptoms.

## Introduction

The rodent parafascicular nucleus (Pf) of the thalamus provides glutamatergic projections to the basal ganglia (BG), a set of subcortical nuclei critical for action selection ^1, 2^. Recent studies have attempted to elucidate the functional significance of Pf interactions with the BG, with emphasis on its thalamostriatal projections ^2–4^, implicating this projection in learning and behavioral flexibility ^5–7^. Furthermore, connectivity with numerous BG nuclei has also made the Pf an exploratory target for deep brain stimulation (DBS) in Parkinson’s disease (PD) patients. Of particular interest are Pf projections to the subthalamic nucleus (STN): the most common target for DBS in PD ^8, 9^. Yet little is known about how the Pf contributes to movement and the functional significance of Pf-STN projections remains unknown.

In this study, we investigated the functional significance of Pf projections to the STN, specifically its contribution to movement. We found that neurons in the Pf represent velocity during self-initiated movements. Moreover, cell-type and pathway specific optogenetic excitation of Pf glutamatergic neurons projecting to the STN generated movements, whereas Pf thalamostriatal projections did not. Optogenetic excitation of the Pf-STN pathway in bilateral 6-hydroxydopamine (6-OHDA) PD mice with complete akinesia restored locomotion and multiple natural behaviors. Lastly, viral-based retrograde tracing from the Pf and transsynaptic tracing of Pf-defined cells in the STN reveal putative circuits that could mediate our observed Pf-STN PD rescue.

## Results

### Neurons in the Pf Nucleus Represent Velocity

To understand the relationship between Pf neural activity and movement, we wirelessly recorded *in vivo* single-unit activity in the Pf and tracked movements of freely behaving mice with a three-dimensional (3D) motion capture system ^10–12^. Water-deprived mice with chronically implanted electrode arrays in the Pf were trained to track a lever arm for a set interval of time (1 s) to receive a sucrose reward (Figures 1A and 1B). One infrared marker placed on the moving reward arm and two infrared markers placed on each side of a head bar were used to track head movements. This task allows recordings of neural activity during continuous, self-initiated orienting and turning behaviors while mice track a moving target. We found that a subset of neurons in the Pf represent velocity primarily in the ipsiversive direction (27%) relative to the recording hemisphere (Figure 1C, *r*^2^ = 0.99, *n* = 21; Figure 1D, left panel; Figure 1E). These velocity neurons are most active during reward arm tracking in the ipsiversive direction, but are anti-correlated in the contraversive direction (Figure 1D, right panel; Figure 1E). On the other hand, far fewer Pf neurons (5%) were found to represent contraversive velocity (Figure S1). This is the first report of such a robust and continuous relationship with ipsiversive turning velocity anywhere in the brain.

**Figure 1.**
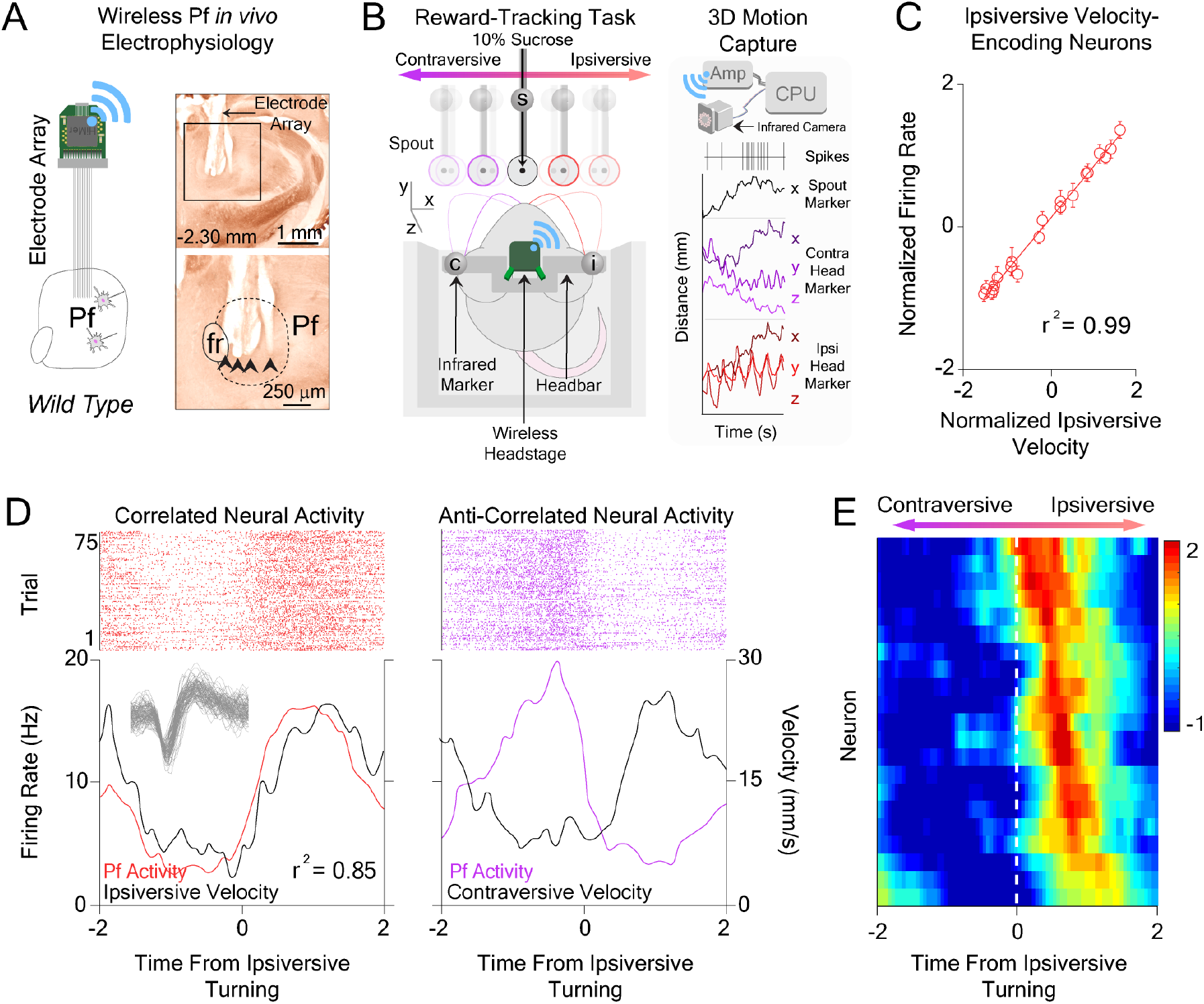
Neural Activity in the Pf Represents velocity. (**A**) Chronically implanted 16-channel electrode array into the Pf of *wild type* mice. Inset shows coronal sections through the thalamus and placement of electrode array into the Pf. Black arrows indicate electrode tips. (**B**) Schematic of reward tracking experimental design. Reflective markers were placed on the head (c and i) of water-deprived freely behaving mice and the reward spout (s) to track movements in relation to a 10% sucrose reward during wireless recordings of Pf neural activity. Right panel illustrates simultaneous neural recordings during acquisition of movement kinematic information from reflective markers in Cartesian space while mice track a reward lever. (**C**) Correlation between normalized Pf firing frequency and normalized ipsiversive velocity across Pf ipsiversive velocity representing neurons (*r*^2^ = 0.99, *n* = 21). (**D**) Representative peri-event histograms showing correlation of Pf neural activity with ipsiversive velocity (left panel) and anti-correlation with contraversive velocity (right panel) for 75 trials. Inset shows Pf waveform. (**E**) Z-scored spike density heat map. Each row represents activity from a single Pf ipsiversive velocity representing neuron with respect to time from ipsiversive turning (*n* = 21). Right side represents ipsiversive direction, left side represents contraversive direction.

### Optogenetic Excitation of Pf Vglut2+ Neurons Initiates Movement

To understand the contribution of the Pf to movement initiation, we optogenetically modulated Pf neural activity while monitoring movement kinematics during 3D motion capture of freely behaving mice in an open-field arena (Figures 2A and 2B). We first performed cell-type nonselective optogenetic excitation of neurons in the Pf by infecting neurons with an adeno-associated virus (AAV) under a ubiquitous neuronal promoter in *wild type* mice (10 mW, 5 ms pulses, AAV5.Syn.Chronos.GFP.WPRE.bGH, *n* = 7, Figure S2A). Unilateral Pf cell-type nonselective optogenetic excitation produces ipsiversive turning and postural changes (Figure S2B). To limit excitation to Pf glutamatergic projection neurons (*Vglut2+*), we injected an AAV with Cre-dependent channelrhodopsin (ChR2) in *Vglut2-ires-Cre* transgenic mice (Figure 2A). *Vglut2* is a selective marker for glutamatergic projection neurons in the thalamus, unlike cortical projection neurons that use *Vglut1* ^13, 14^. We found that cell-type selective excitation of Pf *Vglut2+* neurons produces the same ipsiversive turning and postural changes observed during Pf cell-type nonselective excitation (Figures 2C-E; Movie S1; Vglut2::ChR2^Pf^, *n* = 6; Vglut2::eYFP^Pf^ control mice, *n* = 6). In both cohorts, increasing the excitation frequency increases the speed of ipsiversive turning (Figures 2F and S2B, left panels). Furthermore, a single laser pulse produces a relatively fixed amount of ipsiversive turning, regardless of the frequency of excitation (Figures 2F and S2B, right panels). Bilateral optogenetic excitation slows movement throughout the stimulation period (Figure S3A). These results suggest that Pf activation orients mice in the ipsiversive direction, and bilateral excitation engages antagonistic behaviors.

**Figure 2.**
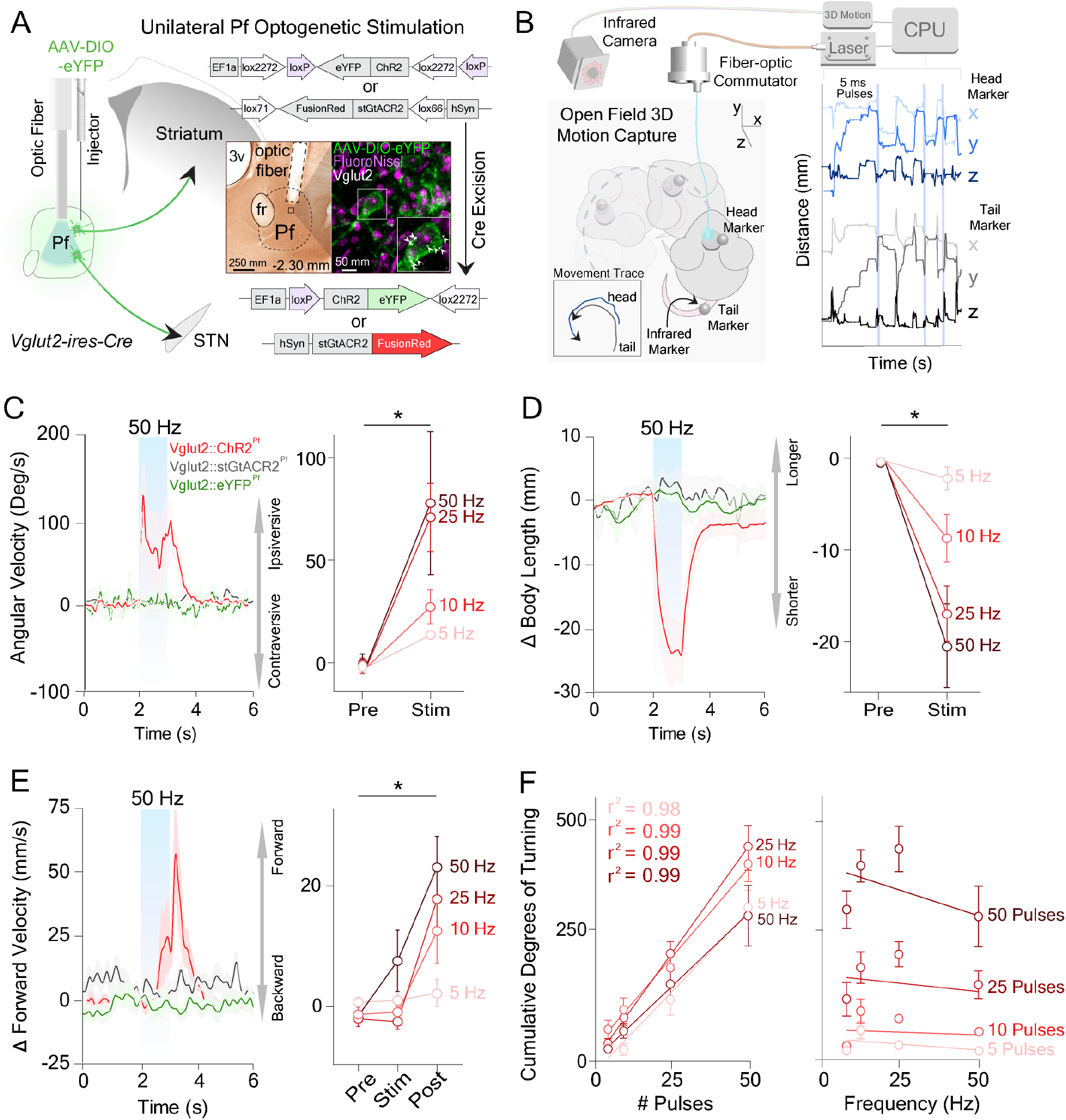
Optogenetic Excitation of Pf *Vglut2+* Neurons Initiates Movement. (**A**) Optogenetic stimulation of glutamatergic neurons in the Pf of *Vglut2-ires-Cre* transgenic mice. Coronal sections through the thalamus showing optic fiber placement above virally infected Pf *Vglut2+* neurons. White arrows indicate colocalization of *Vglut2* (white) in eYFP viral infected cell. (**B**) Open-field 3D motion capture during optogenetic stimulation. Infrared cameras captured the position of reflective markers on the head and tail of freely behaving mice. (**C**) Unilateral optogenetic excitation of Pf *Vglut2+* cell-bodies caused a significant increase in angular velocity (two-way RM ANOVA, pre vs. post stimulation (*F*(1,5) = 12.46, *p* = 0.017, *n* = 6)). (**D**) Unilateral optogenetic excitation of Pf *Vglut2+* cell-bodies caused a significant bending of the body as measured by the distance between the head (h) and tail (t) reflective markers (two-way RM ANOVA, pre vs. post stimulation (*F*(1,5) = 21.59, *p* = 0.0056, *n* = 6)). (**E**) Unilateral optogenetic excitation of Pf *Vglut2+* cell-bodies caused a significant change in forward velocity post-stimulation (two-way RM ANOVA, epoch (*F*(2,10) = 16.84, *p* = 0.006, *n* = 6)). (**F**) Cumulative ipsiversive rotations in degrees significantly increased as a function of the number of excitation pulses (left), regardless of the excitation frequency (right) (Linear regression analyses, all *r*^2^ ≥ .99). Error bars = mean ± SEM.

On the other hand, inhibition of Pf *Vglut2+* neurons with halorhodopsin (rAAV5-EF1α-DIO-eNpHR3.0-eYFP) did no generate turning (Figure S4; Vglut2::eNpHR3.0^Pf^, *n* = 6). Because optogenetic inhibition with eNpHR3.0 might not be sufficient to shut down Pf activity in a normally behaving animal ^15^, we also used soma-targeted *Guillardia theta* anion-conducting ChR2 (AAV1-hSyn-SIO-stGtACR2-FusionRed) to achieve high-efficiency optogenetic silencing of Pf *Vglut2+* neural activity (Vglut2::stGtACR2^Pf^, *n* = 6) ^16^. Similar to stimulating eNpHR3.0 infected Pf neurons, we did not observe any significant effects on turning by stimulating stGtACR2 infected Pf neurons (Figure 2C-E, Figure S4). Interestingly, we did observe a significant lowering of the head in both eNpHR3.0 and stGtACR2 infected animals (Figure S4C). We next tested whether a pharmacological inhibitor of Pf would produce turning by unilaterally injecting muscimol, a GABAA agonist, through chronically implanted guide cannulae (Figure S5; *n* = 8). Unilateral muscimol injections produced contraversive turning in a dose-dependent manner and is consistent with previous observations in rats (Figure S5B and S5C; Movie S2) ^17^. Thus, Pf excitation and inhibition appear to have opposite effects on turning behavior.

### Optogenetic Excitation of Pf Glutamatergic Projections to the STN Produces Movement

Our *in vivo* recording data are in accord with our open-field optogenetic excitation results, suggesting that the Pf contains neurons that could drive ipsiversive turning and orienting behaviors. Two major BG targets of the Pf are the striatum and STN, which are both involved in BG circuits for action initiation (Figure S6). To elucidate the functional contributions of Pf projections to these BG nuclei, we first performed optogenetic excitation of the Pf-striatum pathway. In the first experimental cohort, we injected a Cre-dependent virus containing ChR2 into the Pf of *Vglut2-ires-Cre* transgenic mice and implanted optic fibers above the dorsal striatum (Figure 3A, left panel; Vglut2::ChR2^Pf-striatum^, *n* = 8). We found that unilateral terminal excitation of Pf-striatum neurons does not generate ipsiversive turning or postural changes (Figures 3B-D). We next tested whether direct excitation of cell bodies in the Pf projecting to the striatum would produce movement. To achieve this, we injected AAV-Retro2-Cre, a virus with retrograde access to projection neurons, into the striatum and a Cre-dependent virus containing ChR2 into the Pf of *wild type* mice (Figure 3A, right panel; Retro::ChR2^Pf-striatum^, *n* = 8; Retro::eYFP^Pf-striatum^ control mice, *n* = 6) ^18^. This retrograde strategy allows pathway specific optogenetic excitation, thereby activating a greater proportion of Pf-striatum neurons as compared to Pf-Striatum terminal stimulation. We found that this Pf-striatum cell-body excitation also does not elicit movement (Figures 3B-D).

**Figure 3.**
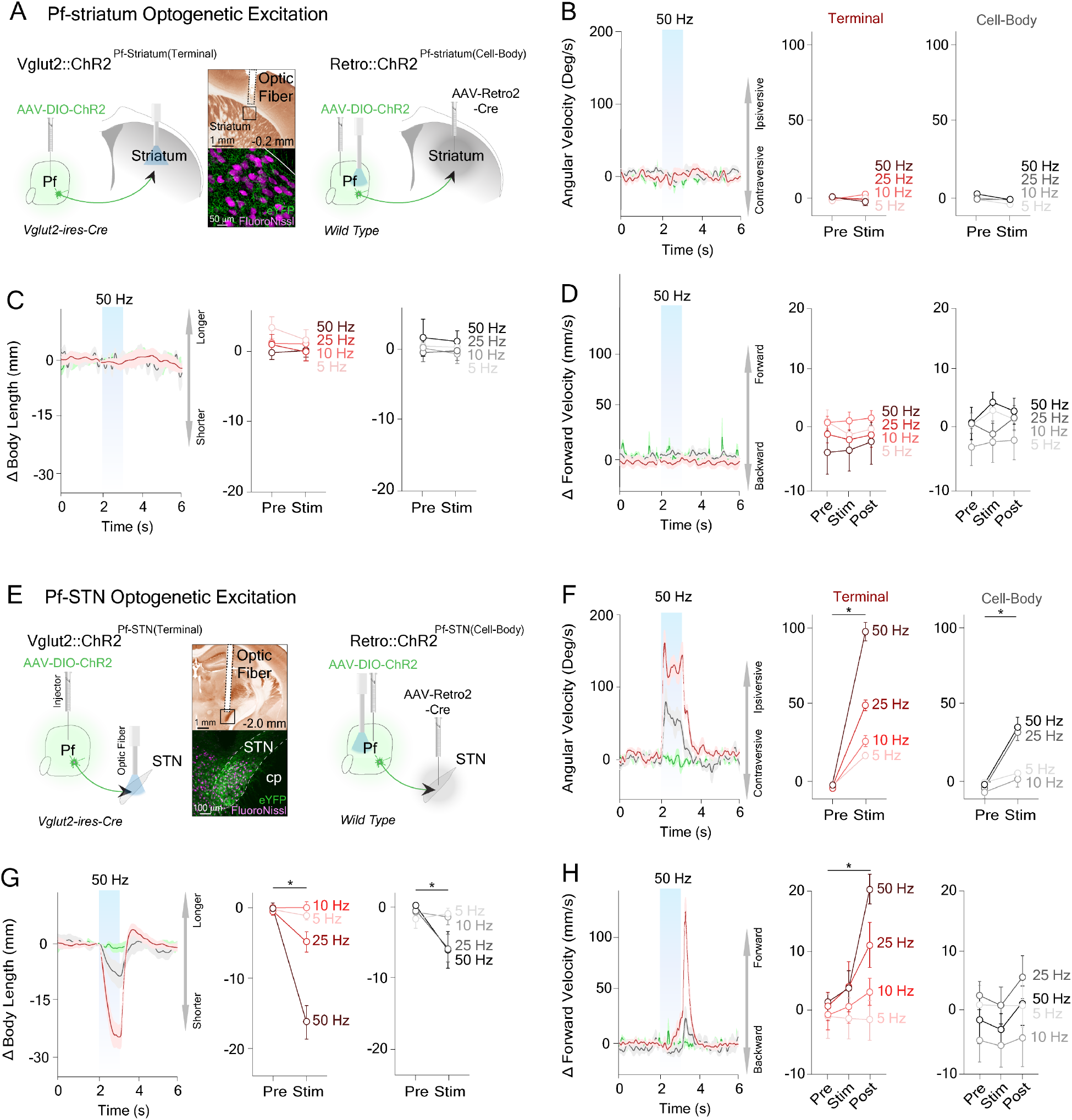
Optogenetic Excitation of the Pf-STN, but not Pf-striatum Pathway Produces Ipsiversive Turning and Postural Changes. (**A**) Pathway specific thalamostriatal (Pf-striatum) optogenetic excitation experimental designs during 3D motion capture. For Pf-striatum terminal excitation experiments, Pf *Vglut2+* infected terminals (AAV-DIO-ChR2) in the striatum were excited with an optic fiber in *Vglut2-ires-Cre* transgenic mice. Coronal section shows optic fiber placement above virally infected Pf-striatum terminals. For Pf-striatum cell-body excitation experiments, infected cell-bodies in the Pf (AAV-DIO-ChR2) projecting to the striatum (AAV-Retro2-Cre) were activated in *wild type* mice. (**B**) No significant changes in angular velocity were observed during Pf-striatum cell-body and terminal excitation (Pf-striatum cell-body, grey trace: two-way RM ANOVA, pre vs. post stimulation (*F*(1,7) = 3.66, *p* = 0.0973, *n* = 8), Pf-striatum terminal, red trace: two-way RM ANOVA, pre vs. post stimulation (*F*(1,7) = 0.4119, *p* = 0.5415, *n* = 8)). Retro::eYFP^Pf-striatum^ control mice shown in green trace (*n* = 6). (**C**) No significant changes in body bending were observed during Pf-striatum cell-body and terminal excitation (Pf-striatum cell-body: two-way RM ANOVA, pre vs. post stimulation *(F*(1,7) = 0.1883, *p* = 0.6774, *n* = 8), Pf-striatum terminal: two-way RM ANOVA, pre vs. post stimulation (*F*(1,7) = 6.836, *p* = 0.0347, *n* = 8)). (**D**) No significant changes in forward velocity were observed during Pf-striatum cell-body and terminal excitation (Pf-striatum cell-body: two-way RM ANOVA, epoch (*F*(2,14) = 1.42, *p* = 0.2745, *n* = 8), Pf-striatum terminal: two-way RM ANOVA, epoch (*F*(2,14) = 0.8859, *p* = 0.4343, *n* = 8)). (**E**) Pathway specific thalamo-subthalamic (Pf-STN) stimulation designs during 3D motion capture. For Pf-STN terminal excitation experiments, Pf *Vglut2+* infected terminals (AAV-DIO-ChR2) in the STN were excited with an optic fiber in *Vglut2-ires-Cre* transgenic mice. Coronal section through the thalamus shows optic fiber placement above virally infected Pf-STN *Vglut2+* terminals. For Pf-STN cell-body excitation experiments, infected cell-bodies in the Pf (AAV-DIO-ChR2) projecting to the STN (AAV-Retro2-Cre) were excited with an optic fiber placed above the Pf in *wild type* mice. (**F**) Optogenetic excitation of Pf-STN cell-bodies and terminals caused a significant increase in angular velocity ipsiversively (Pf-STN cell-body, grey trace: two-way RM ANOVA, pre vs. post stimulation (*F*(1,6) = 26.76, *p* = 0.0021, *n* = 7), Pf-STN terminal, red trace: two-way RM ANOVA, pre vs. post stimulation (*F*(1,7) = 62.24, *p* <0.0001, *n* = 8)). Retro::eYFP^Pf-STN^ control mice shown in green trace (*n* = 6) (**G**) Excitation of Pf-STN cell-bodies and terminals caused a significant bending of the body as measured by the distance between the head (h) and tail (t) reflective markers (Pf-STN cell-body: two-way RM ANOVA, pre vs. post stimulation (*F*(1,6) = 9.326, *p* = 0.0224, *n* = 7), Pf-STN terminal: two-way RM ANOVA, pre vs. post stimulation (*F*(1,7) = 33.51, *p* = 0.0007, *n* = 8)). (**H**) Excitation of Pf-STN cell-bodies and terminals caused a significant increase in forward velocity post-stimulation (Pf-STN cell-body: two-way RM ANOVA, epoch *(F*(2,12) = 15.61, *p* = 0.0005, *n* = 7), Pf-STN terminal: two-way RM ANOVA, epoch (*F*(2,14) = 32.25, *p* <0.0001, *n* = 8)). Error bars = mean ± SEM. Abbreviations – cp, cerebral peduncle.

We next tested whether excitation of Pf projections to the STN would produce ipsiversive turning and postural changes. We found that optogenetic excitation of Pf terminals in the STN produces ipsiversive turning (Figures 3E-H; Figure S7; Movie S3; Vglut2::ChR2^Pf-STN^, *n* = 8). These elicited movement effects during Pf-STN terminal optogenetic excitation are similar to those observed during Pf *Vglut2+* cell-body excitation (Figure 2). We replicated our Pf-STN optogenetic terminal excitation results by exciting cell-bodies of Pf neurons that project to the STN using the retrograde viral strategy previously described (Figure 4E, left panel; Figures 4F-H; Retro::ChR2^Pf-STN^, *n* = 8; Retro::eYFP^Pf-STN^ control mice, *n* = 6). Additionally, we observed that bilateral excitation of the Pf-STN pathway does not slow movement (Figure S3B). Together, these data demonstrate an extrastriatal route by which the thalamus can rapidly influence BG motor output.

**Figure 4.**
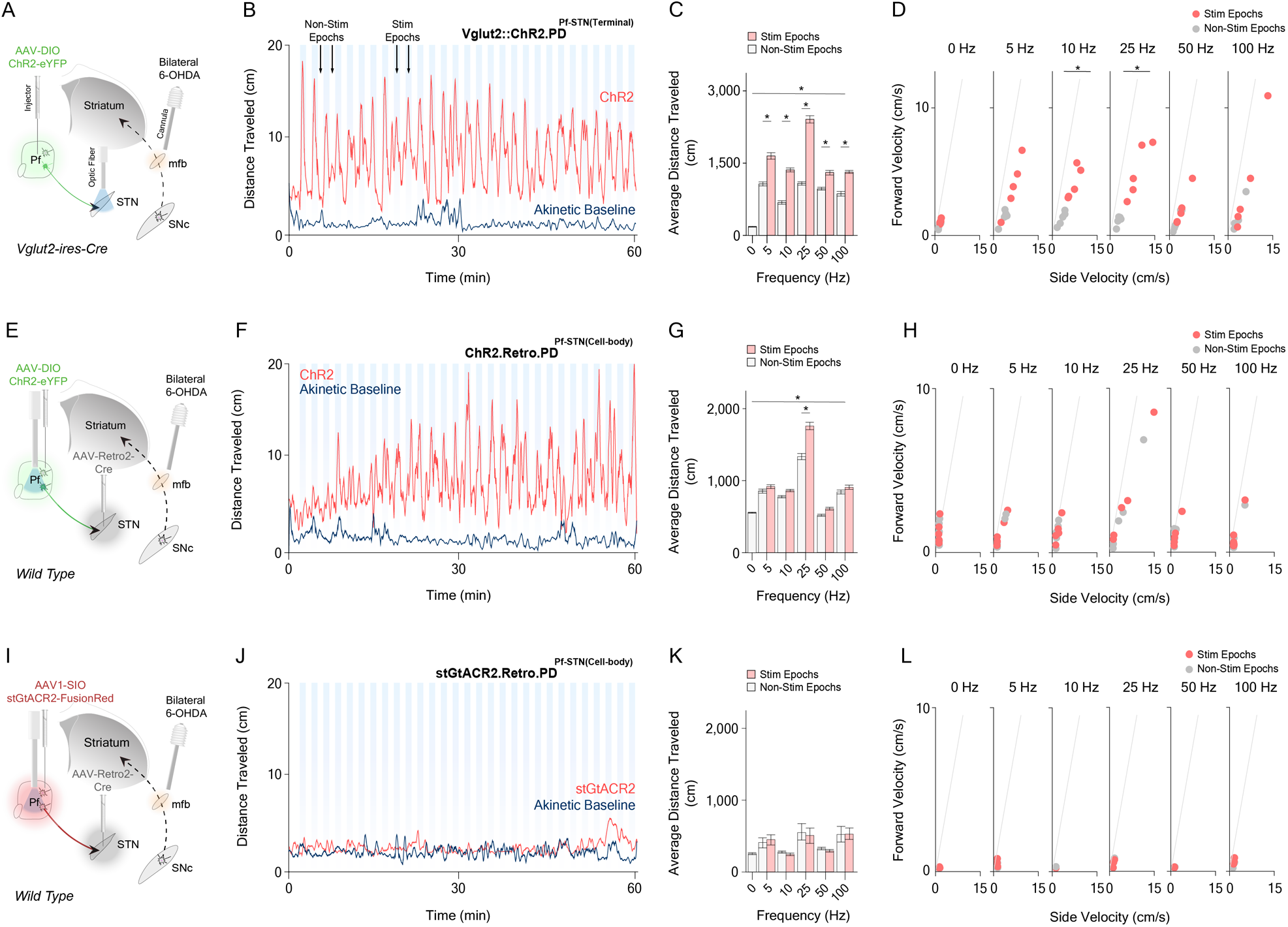
Bilateral Optogenetic Pf-STN Terminal and Cell-body Excitation in Bilateral 6-OHDA Parkinsonian Mice Rescues Akinesia and Promotes Locomotion. (**A**) Pf-STN terminal excitation in bilateral-PD mice. An injection of AAV-DIO-ChR2 into the Pf of *Vglut2-ires-Cre* transgenic mice allowed optogenetic excitation of Pf-STN terminals via an optic fiber implanted above the STN. Bilateral 6-OHDA injections into the mfb through guide cannulae generated a bilateral-PD mouse model four weeks post-surgery. (**B**) Representative example of a 25 Hz activation of PF-STN terminals over 1 hr, showing increased distance traveled across stimulation and non-stimulation epochs. No increase in distance traveled was observed in the akinetic baseline condition. (**C**) Pf-STN terminal excitation in bilateral-PD mice increased the average distance traveled compared to the akinetic baseline condition (two-way RM ANOVA, ChR2 vs. baseline (*F*(2,60) = 0.760.3, *p* <0.0001, *n* = 5)). Post hoc comparison using Tukey’s multiple comparison test indicated that the average distance traveled during all stimulation epochs was significantly higher. (**D**) Average forward and side velocities was increased due to optogenetic excitation. For 10 Hz and 25 Hz conditions, velocities during optogenetic excitation was significantly increased (Hotelling’s t-tests, *p* <0.05 for 10 Hz (*F* = 9.7049, *T* = 22.18) and 25 Hz (*F* = 7.640, *T* = 16.9)). (**E**) Pf-STN cell-body excitation in bilateral-PD mice. An injection of AAV(retro2).hSyn.EF1α.Cre.WPRE into the STN paired with an injection of rAAV5.EF1α.DIO.ChR2.(H134R).eYFP into the Pf of *wild type* mice allowed optogenetic excitation of Pf-STN cell-bodies via an optic fiber implanted above the Pf. (**F**) Representative example of 25 Hz ABA optogenetic stimulation session of Pf-STN cell-bodies over 1 hr, showing increased distance traveled across stimulation and non-stimulation epochs. (**G**) Pf-STN cell-body excitation in bilateral-PD mice significantly increased the average distance traveled compared to the akinetic baseline condition (two-way RM ANOVA, ChR2 vs. baseline (*F*(2,60) = 86.70, *p* <0.0001, *n* = 5)). *Post hoc* comparison using Tukey’s multiple comparison test indicated that the average distance traveled during the 25 Hz stimulation epoch was significantly different its non-stimulation epoch (M = 402.9, SD = 68.09). (**H**) Average forward and side velocities during stimulation and non-stimulation Pf-STN cell-body excitation epochs for each bilateral-PD mouse (*n* = 5). Velocities between epochs were not significantly different (Hotellings t-tests, *p* >0.05). (**I**) Pf-STN cell-body inhibition in bilateral-PD mice. An injection of AAV(retro2).hSyn.EF1α.Cre.WPRE into the STN paired with an injection of AAV1-hSyn-SIO-stGtACR2-FusionRed into the Pf of *wild type* mice allowed optogenetic inhibition of Pf-STN cell-bodies via an optic fiber implanted over the Pf. (**J**) Representative example of 25 Hz ABA optogenetic stimulation session of Pf-STN cell-bodies over 1 hr. Optogenetic Pf-STN cell-body inhibition did not increase the distance traveled across stimulation and non-stimulation epochs. (**K**) Pf-STN cell-body inhibition in bilateral-PD mice did not significantly increase the average distance traveled compared to the akinetic baseline condition (two-way RM ANOVA, ChR2 vs. baseline (*F*(2,60) = 5.09, *p* = 0.4840, *n* = 3)). (**L**) Average forward and side velocities during stimulation and non-stimulation Pf-STN cell-body inhibition epochs for each bilateral-PD mouse (*n* = 3). Velocities between epochs were not significantly different (Hotellings t-tests, *p* >0.05). Error bars = mean ± SEM. Abbreviations – mfb, medial forebrain bundle; SNc, substantia nigra pars compacta.

### Optogenetic Excitation of Pf Projections to the STN Rescues Motor Deficits in a Bilateral 6-OHDA Parkinsonian Mouse Model

Our results showing Pf-STN optogenetic excitation generating movement suggest a circuit mechanism for DBS upstream of the STN. In other words, at least some of the therapeutic effects of conventional STN-DBS may be due to stimulation of the Pf-STN pathway. To test this hypothesis, we used a bilateral 6-OHDA PD mouse model in which dopamine neurons from the substantia nigra pars compacta (SNc) are depleted by injecting 6-OHDA into the medial forebrain bundle (mfb) while modulating Pf-STN pathway activity (Figure 4A) ^19, 20^. Compared to the unilateral model, the bilateral 6-OHDA PD model more closely resembles severe PD, which is not typically restricted to one hemisphere. Because this bilateral-PD model produces severe akinesia, it is rarely used in previous research. However, using this model provides an enhance opportunity to measure behavioral rescue resultant from Pf-STN pathway stimulation. As expected, bilateral 6-OHDA injections produced >80% dopamine depletion in the striatum and nearly complete akinesia in mice (Figure 4B; Figure S8).

We excited Pf-STN terminals using *Vglut2+* transgenic mice while recording movement in an open-field arena (Figure 4A; Figure S9A; Movie S4). An ABA stimulation design was used to perform optogenetic excitation (ChR2) or inhibition (stGtACR2) of Pf-STN cell-bodies in bilateral-PD mice over a 1 hr experimental session (Figure 4B; Figure S9B; Vglut2.PD::ChR2^Pf-STN(terminal)^, *n* = 5). This ABA stimulation design allows measurement of locomotion during stimulation epochs (30 min total) and examination of any residual movement effects during interleaved non-stimulation epochs (30 min total). We show that Pf-STN terminal optogenetic excitation in bilateral-PD mice significantly increases locomotion as measured by the average distance traveled compared to the akinetic baseline condition across all excitation frequencies (Figure 4C). A significant increase in the average distance traveled was also observed between non-stimulation and stimulation epochs across frequencies. We next examined changes in locomotion within velocity space across non-stimulation and stimulation epochs. Stimulation at 10 Hz or 25 Hz increased turning velocity (Figure 4D). Together, these results suggest that Pf *Vglut2+* projections to the STN can be excited to rescue akinesia in bilateral-PD mice.

We next examined whether optogenetic excitation of Pf-STN cell-bodies would produce the same locomotor rescue in bilateral-PD mice. To achieve this, we used the retrograde viral strategy in *wild type* mice as previously described (Figure 4E; Movie S5; Retro.PD::ChR2^Pf-STN(cell–body)^, *n* = 5). Like Pf-STN terminal excitation, we found a significant increase in the average distance traveled compared to the akinetic baseline condition across all excitation frequencies (Figures 4F and 4G). Interestingly, the only significant increase in the average distance traveled between non-stimulation and stimulation epochs was in the 25 Hz condition. No significant change in velocity between non-stimulation and stimulation epochs was observed (Figure 4H). We next wanted to test whether inhibition of Pf projections to the STN could rescue akinesia in bilateral-PD mice (Figure 4I; Retro.PD::stGtACR2^Pf-STN(cell–body)^, *n* = 3). We found no locomotor rescue during Pf-STN cell-body inhibition in bilateral-PD mice (Figures 4J-L). Collectively, these results show that Pf-STN terminal excitation is more efficacious than Pf-STN cell-body excitation at restoring locomotion in bilateral-PD mice.

### Optogenetic Excitation of Pf Projections to the STN Restores Natural Behaviors in a Bilateral 6-OHDA Parkinsonian Mouse Model

Spontaneous behaviors can be highly diverse. While traditional measures only focus on specific aspects of behavior (e.g. average distance travelled or velocity), they do not provide a comprehensive view of different types of behaviors affected by a neural manipulation. To quantify natural behavioral states during our PD rescue experiments, we applied an unsupervised Auto-Regressive Hidden Markov Model (AR-HMM) to our 2D video recordings ^21^. Each video frame (50 f/s) was assigned a behavioral label, which was used to calculate the probability of using each behavioral state, as well as the probabilities of transitioning between states (Figure 5A). The AR-HMM clustered behaviors into kinetic and akinetic groups and revealed a rescue of distinct kinetic behavioral states during both Pf-STN terminal and cell-body optogenetic excitation in bilateral-PD mice (Figure 5; Figure S10A; Movie S6). If we compare the baseline akinetic condition to excitation epochs, excitation restores multiple natural behaviors such as locomotion, turning, perimeter exploration, rearing, and head movement, while reducing immobility behaviors (Hotelling’s t-tests, *p* <0.0001 for all comparisons except 5 Hz (*p* > 0.05)). This behavioral rescue is shown by the higher state probability of kinetic behaviors and lower state probability of akinetic behaviors during optogenetic excitation. In fact, the behavioral state probability during excitation approached that of naïve animals in numerous behaviors (Figure 5B; Figure S10B).

**Figure 5.**
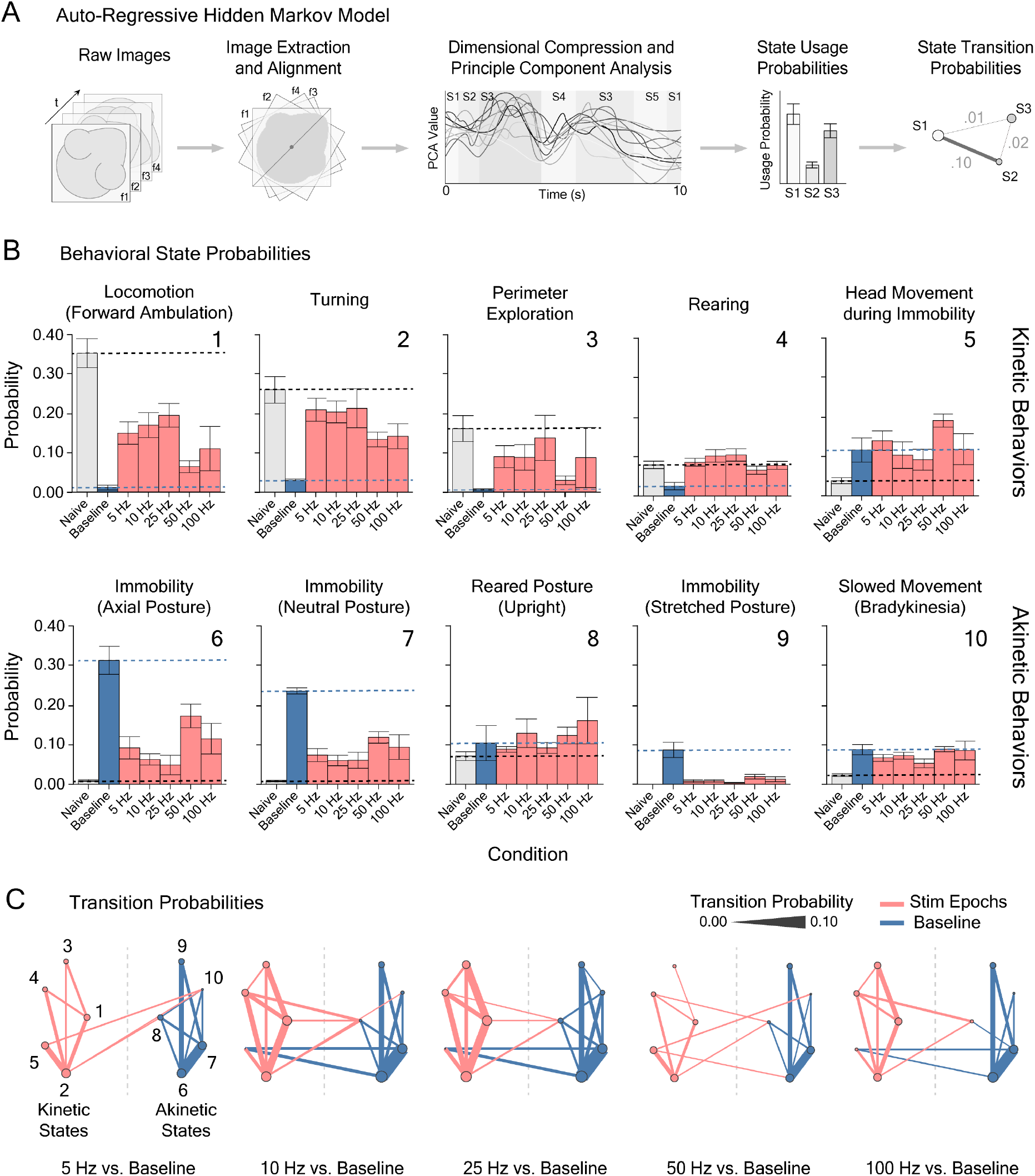
Optogenetic Excitation of Pf-STN Terminals in Bilateral 6-OHDA Parkinsonian Mice Restores Natural Behaviors as Revealed by an AR-HMM. (**A**) Data processing pipeline for an Autoregressive Hidden Markov Model (AR-HMM) on 2D video capture from Pf-STN optogenetic terminal excitation experiments. Abbreviations – f, frame; S, state; t, time. (**B**) Probability of usage of AR-HMM identified kinetic (top row) and akinetic (bottom row) behavioral states during Pf-STN terminal excitation epochs (ChR2, red) across various frequencies (5, 10, 25, 50, and 100 Hz) in bilateral 6-OHDA PD mice. Stimulation groups are statistically different from the akinetic baseline condition (blue) (Hotelling’s t-tests, *p* <0.0001 for all comparisons except 5 Hz (*p* > 0.05)). Error bars = SEM. Dotted lines represent mean in akinetic baseline and naïve conditions. Numbers represent behavioral states shown in (C). (**C**) Pf-STN terminal excitation increases mobility between kinetic behavioral states compared to the akinetic baseline condition in bilateral-PD mice. Bigrams illustrating transition probabilities between AR-HMM identified kinetic (left) and akinetic (right) states across frequencies during Pf-STN terminal excitation epochs (red) versus the akinetic baseline condition (blue). Circle diameter represents state probability. Line thickness represents transition probability. Numbers represent behavioral states identified in (B): 1) locomotion (forward ambulation); 2) turning; 3) perimeter exploration; 4) rearing; 5) head movement during immobility; 6) immobility (axial posture); 7) immobility (neutral posture); 8) reared posture; 9) immobility (stretched posture); 9) slowed movement (bradykinesia). Error bars = mean ± SEM.

**Figure 6.**
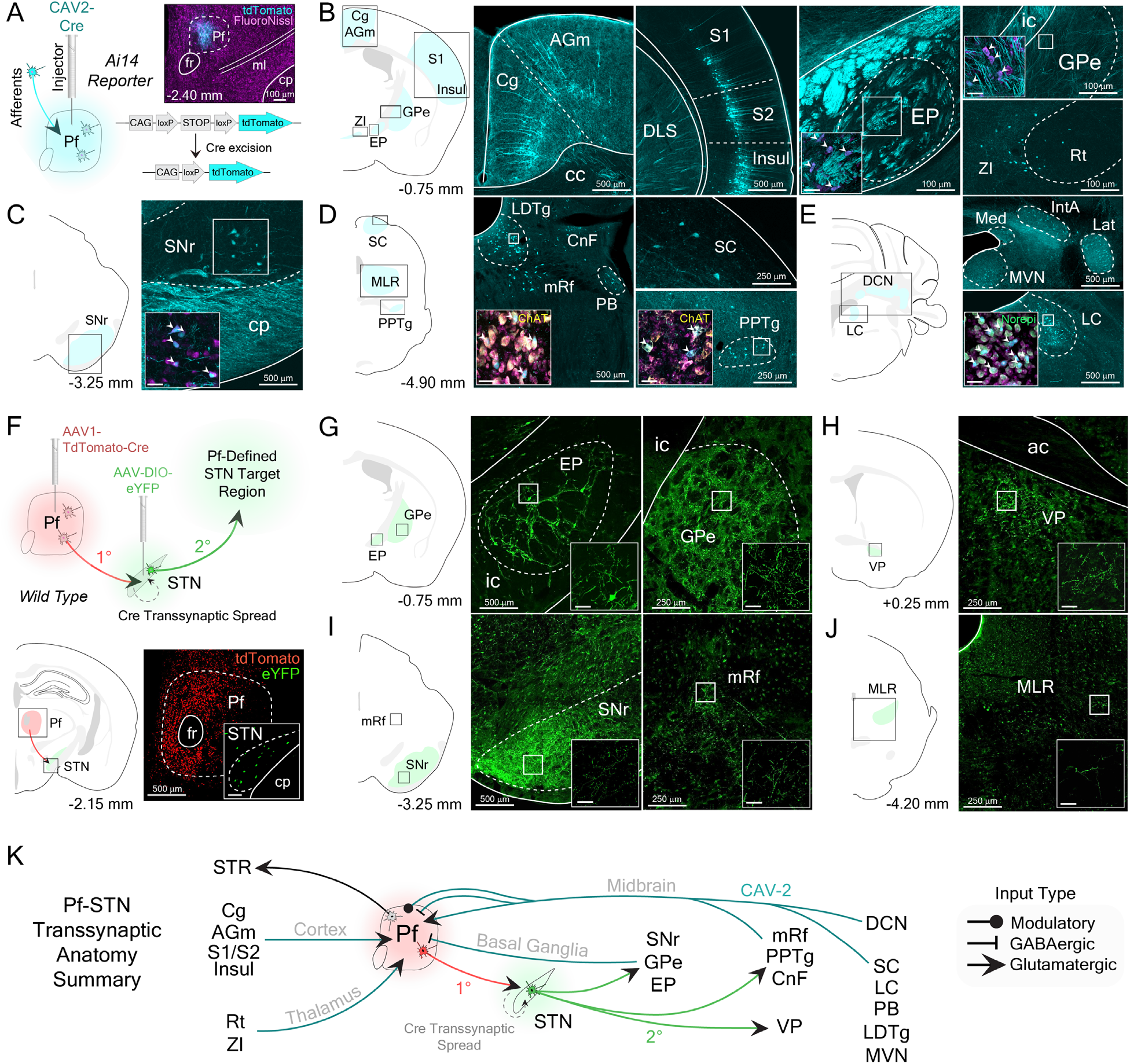
Pf-Defined Projections through the STN Transsynaptically Target Output Nuclei of the Basal Ganglia and Brainstem Locomotor Regions. (**A**) Representative example of a CAV2-Cre injection into the Pf of an *Ai14* transgenic mouse to visualize afferent projections. Neurons in *Ai14* mice fluoresce (tdTomato, cyan) when infected in the presence of a Cre-recombinase virus. Inset shows CAV2-Cre injection site into the Pf. (**B-E**) Retrograde labeled cells from Pf CAV2-Cre injection shown in panel A; AGm, Cg, EP (rodent homolog of primate GPi), GPe, Insul, Rt, S1, S2 and ZI (B); SNr (C); CnF, LDTg, mRf, PB, PPTg, and SC (D); IntA, Lat, LC, Med, and MVN (E). Insets show retrograde labeled cells colocalized with neuromodulatory cell-type markers (ChAT, Norepi). Scale bars = 100 μm. (**F**) Representative example of Pf-defined transsynaptic projections through the STN. Injection of AAV1.hSyn.tdTomato.Cre into Pf (first order neuron, 1**°**) allows Cre transsynaptic spread into the STN. AAV-DIO-eYFP was subsequently injected into the STN (second order neuron, 2**°**) to visualize Pf-defined subthalamic projections. Coronal schematic shows Pf injection site (tdTomato). Inset shows STN injection site (eYFP). Scale bar = 100 μm. (**G-J**) Coronal sections show terminal labeling sites of Pf-defined subthalamic targets in the EP (rodent homolog of primate GPi) and GPe (G), VP (H), mRt and SNr (I), and CnF (J). Scale bar = 100 μm. (**K**) Summary of viral-based retrograde Pf and transsynaptic Pf-STN anatomy results. The Pf receives glutamatergic, GABAergic, and modulatory input from mesencephalic brain regions. The Pf also receives GABAergic feedback from Pf-STN transsynaptic targeted basal ganglia nuclei, in addition to glutamatergic input from cortex. Abbreviations – ac, anterior commissure; AGm, agranular medial cortex; cc, corpus callosum; Cg, cingulate cortex; ChAT; choline acetyltransferase; cp, cerebral peduncle; CnF, cuneiform nucleus; DCN, deep cerebellar nuclei; DLS, dorsolateral striatum; EP, entopeduncular nucleus; GPe, external segment of the globus pallidus; GPi, internal segment of the globus pallidus; fr, fasciculus retroflexus; Insul, insular cortex; IntA, interposed anterior nucleus of the cerebellum; Lat, lateral cerebellar nucleus; LC, locus coeruleus; LDTg, laterodorsal tegmental nucleus; M1, primary motor cortex; M2, secondary motor cortex; Med, medial cerebellar nucleus; ml, medial lemniscus; MLR, midbrain locomotor region; MLV, medial vestibular nucleus; mRf, mesencephalic reticular formation; NAc, nucleus accumbens; Norepi, norepinephrine; PB; parabrachial nucleus; PPTg, pedunculopontine tegmental nucleus; Rt, reticular nucleus; S1, primary sensory cortex; S2, secondary sensory cortex; SC, superior colliculus; SNr, substantia nigra pars reticulata; STR, striatum; ZI, zona incerta.

We next investigated whether animals during excitation increase their probability of transition amongst kinetic behavioral states. We found that Pf-STN pathway optogenetic excitation in bilateral-PD mice increased the probability of mice transitioning between kinetic behavioral states compared to the akinetic baseline condition, where a high probability of transitioning between akinetic behavioral states is observed (Figure 5C; Figure S10B and S11). The most robust increase in transition probabilities between kinetic states is observed in the 25 Hz condition. Collectively, these data demonstrate that Pf-STN pathway excitation reduces PD motor deficits and increases the occurrence of, and the probability of transition between kinetic behaviors.

### Pf-defined Projections through the STN Transsynaptically Target Output Nuclei of the Basal Ganglia and Brainstem Locomotor Regions

To reveal the potential circuitry responsible for our Pf-STN PD rescue, we first visualized inputs to the Pf using viral-based retrograde tracing by injecting canine adenovirus type 2 with Cre recombinase (CAV2-Cre) into the Pf of *Ai14* reporter mice (Figure 6A). Our CAV2-Cre retrograde tracing shows strong inputs from midbrain neuromodulatory and reticular activating areas, such as the laterodorsal tegmental nucleus, pedunculopontine tegmental nucleus, medial reticular formation, and locus coeruleus (Figures 6B-E). Numerous brain regions implicated in pain and salience detection also project to the Pf, including the superior colliculus, cuneiform nucleus, parabrachial nucleus, and cingulate cortex. Pf afferents were also seen from sensorimotor brain regions (primary motor and sensory cortices, zona incerta, deep cerebellar nuclei) and nuclei of the basal ganglia (entopeduncular nucleus, external globus pallidus, substantia nigra pars reticulata). This is the first comprehensive visualization of Pf afferents in mice and illustrates that the Pf is anatomically situated at the interface between brain regions implicated in arousal and orienting.

We next elucidated downstream circuits that can be affected by Pf activation through the STN, by uncovering the downstream targets of STN neurons that receive synaptic contacts from Pf neurons using a viral-based transsynaptic tracing technology ^22^. To visualize these Pf-defined STN projections, we first injected AAV1.hSyn.tdTomato.Cre into the Pf of *wild type* mice (Figure 6F). This virus enables anterograde transsynaptic spread of Cre into postsynaptically targeted neurons. We then injected a Cre-dependent virus (rAAV5-EF1α-DIO-eYFP) into the STN to visualize Pf-defined STN projections. In other words, only STN neurons that are postsynaptic targets of the Pf will express eYFP. We found that Pf-defined STN efferents terminate in various BG output nuclei, including the substantia nigra pars reticulata (SNr), the external globus pallidus, the entopeduncular nucleus (the rodent homologue of the internal globus pallidus), as well as other key brainstem regions for motor control (Figures 6G-J). This finding shows that, independent of the striatum, the Pf can influence BG output nuclei and brainstem locomotor regions through the STN (Figure 6K). These regions, in turn, also project to the Pf. Furthermore, these results are in agreement with previous observations that activation of BG output nuclei, such as the SNr, produces ipsiversive turning and mediates posture control^23, 24^.

## Discussion

This study elucidated a thalamo-subthalamic pathway regulating movement initiation and revealed a glutamatergic circuit mechanism between the Pf and STN that could partly explain the clinical efficacy of DBS for relief of PD motor symptoms. We found that neurons in the Pf represent velocity during self-initiated movements primarily in the ipsiversive direction, and optogenetic excitation of Pf-STN *Vglut2+* neurons produces ipsiversive turning and postural changes. Surprisingly, Pf-striatum projections are not critical for rapid movement generation. It is possible that Pf projections to the striatum could contribute to higher functions, such as decision making and behavioral flexibility, but do not immediately generate movement ^2, 25^.

Because the STN is the canonical target for DBS in PD patients, we used a common mouse model of PD to test whether selective optogenetic excitation of the Pf-STN pathway could rescue PD-induced motor deficits. We found that excitation of the Pf-STN pathway in bilateral 6-OHDA PD mice restores normal behavior (Figures 4 and 5). To quantify the effect of stimulation on natural behaviors, we used an unsupervised behavioral classification program (AR-HMM), which revealed that Pf-STN excitation in PD mice increased the probability of multiple normal behaviors. We then revealed putative circuits through the BG and locomotor brainstem regions that could mediate our observed Pf-STN PD rescue using viral-based neuroanatomical tracing (Figure 6).

Our findings show that the Pf is a part of a circuit for orienting and steering in mice. Given the strong afferents from neuromodulatory and sensorimotor brain regions, the Pf is anatomically positioned to *combine* somatosensory and proprioceptive information to form representations of movement kinematics. In support of this conclusion, we identified a subset of Pf neurons that are highly correlated with turning velocity, in agreement with our optogenetic excitation results (Figures 1-3). Moreover, we identified downstream locomotor brain regions that could mediate the movement effects observed during Pf-STN optogenetic excitation with viral-based transsynaptic tracing (Figure 6). We thus uncovered a key node in the neural network for left-right steering, with the Pf from each hemisphere responsible for ipsiversive movements.

Bilateral Pf excitation resulted in slowed movement (Figure S3A; Figure S12). This is likely due to simultaneous activation of antagonistic units (e.g., leftward and rightward) from the two hemispheres. It is possible that recruitment of both Pf-STN and other Pf efferents, such as Pf thalamostriatal neurons, produce a conflict at longer stimulation durations and does not mimic the natural activation pattern of these circuits. However, we did not observe this phenomenon during bilateral Pf-STN excitation (Figure S3B). The difference between our Pf-STN and Pf-striatum movement effects cannot simply be explained by differences in penetrance or size of the target region, as the thalamostriatal projections are more massive and target a larger area (Figure 3; Figure S6) ^26^. If anything, this would predict a larger effect of thalamostriatal activation, but the opposite was observed. More importantly, it is well established that striatal activation produces contraversive rather than ipsiversive movement ^27^. Our observation of ipsiversive turning by optogenetic excitation of the Pf-STN pathway therefore supports an extra-striatal route by which Pf influences movement.

With conventional DBS, it is impossible to determine whether the therapeutic effect is due to activating local neural populations, presynaptic axon terminals, or fibers of passage. Nor is it entirely clear whether DBS excites or inhibits local neural elements. As STN-DBS ameliorates key motor symptoms of PD, it is possible that a component of STN-DBS may arise from exciting Pf input to the STN. Our study supports this hypothesis by demonstrating rescue of akinesia in bilateral-PD mice using selective targeting of the Pf-STN pathway (Figure 4). Previous work showed that optogenetic excitation of STN neurons did not rescue akinesia, but that inputs from motor cortex were more important ^28^. These results suggest the possibility that conventional DBS produces antidromic activation of neurons presynaptic to the STN. Here we show that the Pf is a key region upstream of the STN that could be responsible for DBS rescue of PD symptoms. By exciting thalamic projections to the STN, we were able to restore natural movements in bilateral-PD mice (Figure 5; Figure S10). Thus the therapeutic effects of STN-DBS can be attributed to activating Pf glutamatergic afferents to the STN. This possibility is also supported by a recent study demonstrating increased locomotion in dopamine-depleted mice by optogenetically exciting parvalbumin-positive neurons in the GPe ^20^. Interestingly, these GPe neurons project to the Pf ^29^, suggesting a potential circuit that can be recruited to generate movement in PD mice.

We were able to restore and increase the transition probability of numerous natural behavioral states that are difficult to quantify with conventional measures. For instance, Pf-STN excitation in bilateral-PD mice alleviates numerous axial symptoms, such as gait immobility, and promotes natural postural rearing: symptoms that are not satisfactorily treated with conventional DBS approaches and are often resistant to medication in humans ^8, 30–32^. It is therefore possible that exciting Pf input to the STN in PD patients could abolish pathological STN activity, thus reducing akinesia and promoting volitional movements ^33, 34^.

Interestingly, Pf-STN terminal excitation was more effective at alleviating akinesia and promoting kinetic behaviors in bilateral-PD mice than Pf-STN cell-body excitation (Figures 3 and 4; Figures S10). Similarly, bilateral CM/Pf-DBS in PD patients has limited therapeutic benefits compared to STN-DBS, suggesting that it cannot control PD motor symptoms adequately ^35–37^. Given our finding that non-specific optogenetic excitation slows movement (Figure S3A), it is possible that CM/Pf-DBS in PD patients might recruit conflicting downstream neural circuits that attenuates its therapeutic potential. In contrast, we found that bilateral Pf-STN excitation does not impair movement (Figure S3B). Thus, the benefits derived from STN-DBS in PD patients could be produced by antidromic excitation of Pf input into the STN, bypassing the activation of conflicting upstream neural circuits.

Collectively, our results suggest that Pf-STN projections represent a diencephalic analogue of the hyperdirect pathway, or ‘superdirect’ pathway, allowing the Pf direct access to BG output nuclei and other brainstem locomotor regions critical for movement initiation ^38^. Pf-STN projections may therefore serve as a rapid orienting mechanism to shift the body towards salient stimuli, while Pf thalamostriatal projections can influence the selection of the next motor program. In PD, these projections can be excited to reduce PD motor deficits by putatively recruiting brainstem locomotor regions downstream of the STN.

## Acknowledgements

We thank Fengxia Allen, Erin Gaidis, Xiaoran Li, Namsoo Kim, and Murray Wickwire for technical assistance, Eric Monson for helpful comments on data representation, and Michael Tadross and Nicole Calakos for helpful comments on the manuscript. This work was supported by grants NS094754 and MH112883 awarded to H.H.Y.

## Author Contributions

Conceptualization, G.D.R.W. and H.H.Y.; Methodology, G.D.R.W., E.A.P., and H.H.Y.; Software, G.D.R.W., R.N.H., and E.A.P; Validation, G.D.R.W. and R.N.H.; Formal Analysis; G.D.R.W., R.N.H., and E.A.P.; Investigation, G.D.R.W. and R.N.H.; Resources; H.H.Y.; Data Curation, G.D.R.W., R.N.H., and E.A.P.; Writing – Original Draft, G.D.R.W.; Writing – Review & Editing, G.D.R.W. R.N.H., E.A.P., and H.H.Y.; Visualization, G.D.R.W.; Supervision, G.D.R.W. and H.H.Y.; Funding Acquisition; H.H.Y.

## Materials and Methods

All experimental procedures were conducted in accordance with standard ethical guidelines and were approved by the Duke University Institutional Animal Care and Use Committee.

### Contact for Reagents and Resource Sharing

The data that support the findings of this study are available from the corresponding author upon reasonable request, Henry H. Yin.

### Experimental and Subject Details

All behavioral data were collected from *wild type* (C57BL/6J, Jackson labs) and *Vglut2-ires-Cre* mice (Slc17a6^tm2(cre)Lowl^, Jackson labs). *Vglut2-ires-Cre* mice have Cre-recombinase expression under the control of *Vglut2* receptor regulatory elements without disrupting endogenous vesicular glutamate transporter 2 expression. Optogenetic control of Pf glutamatergic neurons was achieved with a double-floxed inverted recombinant AAV5 virus injection to express either the excitatory opsin ChR2-eYFP, or the inhibitory opsins eNpHR3.0-eYFP and stGtACR2-FusionRed in *Vglut2+* Pf neurons. Viral infection in the Pf was histologically verified with eYFP or FusionRed imaging colocalized against a Vglut2 antibody and DAPI staining (Figure 1A). *Wild type* mice were used for Pf muscimol inactivation, pathway-specific retrograde Cre expression, and retrograde and transsynaptic tracing experiments. Retrograde anatomical tracing data was collected from Ai14 reporter mice (129S6-Gt(ROSA)26Sor^tm14^(CAG–tdTomato)^Hze^/J, Jackson labs). Ai14 reporter mice harbor a loxP-flanked STOP cassette that is excised in the presence of Cre to promote transcription of a CAG promoter-driven red fluorescent protein variant (tdTomato). A high titer was used (∼3.0 x 10^13^ gc/ml) was used to visualized input-output connectivity. For PD experiments, *Vglut2-ires-Cre* mice were used for Pf-STN terminal excitation experiments and *wild type* mice were used for Pf-STN cell-body excitation experiments.

### Viral Constructs

CAV2-Cre was obtained from Institut de Génétique Moléculaire de Montpellier. AAV5.Syn.Chronos.GFP.WPRE.bGH was obtained from the University of Pennsylvania Vector Core. rAAV5.EF1α.DIO.hChR2(H134R).eYFP, rAAV5.EF1α.DIO.eNpHR3.0.eYFP, rAAV5.EF1α.DIO.eYFP, AAV1.hSyn.tdTomato.Cre, AAV(retro2).hSyn.EF1α.Cre.WPRE, and AAV1-hSyn-SIO-stGtACR2-FusionRed were obtained from the Duke University Vector Core.

### Surgery

Mice were anesthetized with 2.0 to 3.0% isoflurane mixed with 0.60 L/min of oxygen for surgical procedures and placed into a stereotactic frame (David Kopf Instruments, Tujunga, CA). Meloxicam (2 mg/kg) and bupivacaine (0.20 mL) were administered prior to incision. To optogenetically interrogate Pf during open-field experimentation, adult *Vglut2-ires-Cre* mice were randomly assigned to Vglut2::ChR2^Pf^ (*n* = 4, 2 males, 2 females, 8-10 weeks old), Vglut2::eNpHR3.0^Pf^ (*n* = 4, 2 males, 2 females, 8-10 weeks old), Vglut2::stGtACR2^Pf^ (*n* = 3, 1 male, 2 females, 8-10 weeks old), or Vglut2::eYFP^Pf^ groups (*n* = 4, 2 males, 2 females, 8-10 weeks old). Craniotomies were made bilaterally above the Pf and virus was microinjected through a pulled glass pipette at various penetrations and depths (0.6 μL each hemisphere, AP: - 2.10 – 2.50 mm relative to bregma, ML: ± 0.60 – 0.75 mm relative to bregma, DV: −3.70 – 3.10 mm from skull surface) using a microinjector (Nanoject 3000, Drummond Scientific). For pathway specific open field experiments, *wild type* mice were used to selectively target Pf-striatum (*n* = 4, 2 males, 2 females, 8-10 weeks old) and Pf-STN (*n* = 4, 2 males, 2 females, 8-10 weeks old) neurons by bilaterally injecting AAV(retro2).hSyn.EF1α.Cre.WPRE into the entire rostrocaudal and mediolateral extent of the striatum (1 μL each hemisphere, AP: +1.35 – −0.75 mm relative to bregma, ML: ± 1 – 2.75 mm relative to bregma, DV: 2.0 – 3.50 mm from skull surface) or the STN (AP: −1.90 – −2.20 mm relative to bregma, ML: ± 1.40 – 1.80 mm relative to bregma, DV: 4.10 – 4.40 mm from skull surface) in parallel with a Cre-dependent ChR2 virus injection into the Pf. For Pf cell-body stimulation, custom-made optic fibers (5-6 mm length below ferrule, >80% transmittance, 105-mm core diameter) were implanted directly above the nucleus (AP: −2.30 mm relative to bregma, ML: ± 1.20 – 1.30 mm relative to bregma, DV: 2.70 – 2.80 mm from skull surface, 10°). Pf-STN terminals were targeted with optic fibers implanted directly above the STN (AP: −1.80 – 2.0 relative to bregma, ML: ± 1.50 – 1.60 mm relative to bregma, DV: 4.20 mm from skull surface). Pf-striatum terminals were targeted with optic fibers implanted directly above the dorsal striatum (AP: 0.25 – −0.50 mm relative to bregma, ML: ± 2.50 – 2.90 mm relative to bregma, DV: 1.75 mm from skull surface). Pf-STN (*n* = 6, 3 males, 3 females, 10-12 weeks old) and Pf-striatum (*n* = 6, 2 males, 4 females, 10-12 weeks old) controls were generated using the retrograde access viral technique previously described paired with an injection of rAAV5.EF1α.DIO.eYFP into and fiber implants above the Pf. For retrograde anatomy experiments (*n* = 3, 2 males, 1 female, 8-10 weeks old), 15-20nL of CAV2-Cre was injected into the Pf at one depth to prevent leakage (AP: −2.30 mm relative to bregma, ML: ± 0.65 – 0.75 mm relative to bregma, DV: 3.60 – 3.30 mm from skull surface). For transsynaptic tracing experiments (*n* = 3, 1 male, 2 females, 8-10 weeks old), AAV1.hSyn.tdTomato.Cre was injected into the Pf and rAAV5-EF1α-DIO-eYFP was injected into the STN using the coordinates previously listed. For all virus injections, the pipette sat at the last injection depth of each penetration for 20 minutes before being withdrawn from the brain to facilitate uptake. To pharmacologically inactivate the Pf nucleus, muscimol was injected in adult *wild type* mice (*n* = 5, 2 males, 3 females, 10-12 weeks old) through 15-gauge (5 mm length) guide cannulae (Plastics One) positioned above Pf at an angle (AP: −2.30 mm relative to bregma, ML: ± 1.20 - 1.30 mm relative to bregma, DV: 2.60 - 2.80 mm from skull surface, 25°). Bilateral-PD mice (*n* = 13, 7 males, 6 females, 8-10 weeks old) were generated by implanting 15-gauge (7 mm length) guide cannulae implanted above the mfb (AP: −0.95 mm relative to bregma, ML: ± 3.10 mm relative to bregma, DV: 4.10 mm from skull surface, 25°). For electrophysiology experiments (*n* = 6, 3 males, 3 females, 8-10 weeks old), a prefabricated 4×4 electrode array (Innovative Neurophysiology, 150 µm electrode and row spacing) was lowered into the Pf (AP: −2.30 mm relative to bregma, ML: ± 0.65 mm relative to bregma, DV: 3.30 mm from skull surface) at a rate of 300 µm/min. All fibers and cannulae were secured in place with dental acrylic adhered to skull screws. Mice were group housed and allowed to recover for one week before experimentation.

### Optogenetic Stimulation and 3D Motion Tracking

For 3D motion capture experiments, mice were connected to a 473-nm DPSS laser (Shanghai Laser) for optogenetic excitation (ChR2) and inhibition (stGtACR2) via fiber optic cables (105/22A, Precision Fiber) in a square open field arena (22” L, 22” W, 1” H) elevated 3’. Mice were connected to a 589-nm DPSS laser for eNpHR3.0 optogenetic inhibition experiments. The output from each sheathed optic fiber tip was measured (PM120VA, ThorLabs) before each experimental session to obtain a power between 9-10 mW (i.e. ∼8mW power delivered to the stimulation site with a transmittance of ∼85%). A Matlab program interfaced to a National Instruments Box triggered a 5 ms square pulse at varying frequencies (5 - 50 Hz) and durations (50 ms - 20 s). Movements at a millimeter spatial resolution were captured in a Cartesian plane with eight Raptor-H digital infrared cameras (Motion Analysis, CA, 100 Hz sampling rate). The cameras were placed equidistantly around the arena where two infrared spherical markers (6.35 mm diameter) that were located on both the fiber sleeve and tail of the mouse could be recorded. Cameras were calibrated before each experimental session. The output of the laser was channeled through an optic patch cable connected to a commutator above the open field arena. A rotating optical commutator (Doric) divided the beam (50:50) permitting bilateral stimulation. Stimulation was triggered randomly during a 12 to 15 second interval after cessation of the previous stimulation trial using a custom Matlab script to prevent mice from predicting the stimulation onset. Stimulation parameters were consistent within a session, but the order of stimulation (i.e., frequency and duration) was semi-randomized between mice. Unilateral and bilateral stimulation data were taken from the same animals. For unilateral experiments, separate stimulation data sessions were acquired from each hemisphere of the same animal on separate days.

### Pf Pharmacological Inactivation and Medial Forebrain Bundle Lesion

Pf muscimol inactivation experiments were conducted on *wild type* mice 10 days after cannulation surgery. Before each muscimol injection, a 10-minute baseline video was taken. All mice were anesthetized with 2.5% isoflurane and a 33-gauge injector was inserted through the cannula that extended 500 µm beyond the cannula tip. Varying dosages of muscimol (0.1, 0.5, and 1.0 µg/µL) or vehicle (0.9% saline) were unilaterally injected using a custom microinjector and microinfusion pump (PHD 2000, Harvard Apparatus) at a rate of 100 nL/min for 2 min. The injector was withdrawn after 10 minutes to allow sufficient uptake of the drug. Mice were then placed in the center of a rectangular open field arena (17.5” L, 9.5” W, 6.25” H) before recording video of movement 15 min post-injection. An open source software program (Bonsai) using a custom script tracked movements for 10 min ^39^. Two-dimensional coordinates based on the center of mass of each mouse were collected at 30 f/s. Because muscimol has a relatively short half-life (∼ 6 hrs), dosage response experiments were conducted 24 hrs apart in a randomized fashion, resulting in a total of 4 injections per hemisphere for each mouse. The same injection procedures were followed to generate bilateral-PD mice using 6-OHDA (2.0 mg, 1300 nL/hemisphere). A baseline video (50 f/s) was taken 3 days after 6-OHDA injections in a rectangular open field arena (19” L, 10” W, 12” H). An ABA stimulation design was then used to optogenetically excite Pf-STN neurons in bilateral-PD mice across various frequencies (5, 10, 25, 50, and 100 Hz). To achieve short latency control of the laser, we used two Arduinos. The first Arduino served as a data acquisition device to receive digital output signals from the computer. The second Arduino modulated frequency output from the laser using TTL pulses. Stimulation conditions and equipment were the same as previously described for 3D tracking experiments. One-minute stimulation and non-stimulation epochs were interleaved in bilateral-PD experiments to comprise a total duration of 30 minutes for each epoch during a session. Mice were allowed to rest 3 hours between stimulation sessions to prevent any movement rescue effect contamination from the previous session.

### 3D Motion Tracking Analysis

A custom Matlab script pre-processed data smoothed at 6 Hz from Cortex 3D movement tracking acquisition software (Motion Analysis) to output movement kinematic information in Neuroexplorer with respect to stimulation timestamps. Peri-event histograms and rasters were generated in Neuroexplorer and analyzed in GraphPad Prism. Angular Velocity kinematic data was binned at 25 ms and the mean and SEM for each bin across experimental animals was computed to produce movement traces. Kinematic data for head and tail marker distance, head in the z plane, as well as forward velocity were standardized by a Z-score transformation. Angular Velocity was not standardized due to similar velocity profiles across animals. All kinematic statistical analyses were performed on the one second of data preceding stimulation (Pre) compared to the one second of stimulation data (stim).

### Wireless *in vivo* Electrophysiology Recordings

A 128-Channel neural signal data acquisition system (Cerebus, Blackrock Microsystems) recorded action potentials through a miniaturized wireless headstage (Triangle Biosystems) interfaced to manufactured electrode arrays (Innovative Neurophysiology). Data were filtered with both analog and digital bandpass filters before being sampled as previously described ^10^. Collected neural data was sorted offline with OfflineSorter (Plexon) and analyzed in Neuroexplorer (Nexus). Discharges with a signal-to-noise ratio of a least 3:1 were time-stamped at a resolution of 0.1 ms. Waveforms were classified as single units using the following criteria: 1) a signal to noise ratio of at least 3:1; 2) consistent waveforms throughout the recording session; 3) refractory period of at least 800 µs.

### Reward Tracking Task Design and Data Analysis

Pf neural activity was recorded in water deprived mice while mice tracked a moving target controlled by a stepper motor to receive a 10% sucrose reward at 2.7 µL/s (Bipolar, 56.3 x 56.3 mm, DC 1.4A, 2.9 Ω, 1.8 degrees/step, Oriental Motor, USA). The stepper motor was controlled by a custom Matlab script and programmed to traverse a total horizontal distance of 120 mm. One infrared marker placed 20 mm from the sucrose spout and two markers placed on each side of a custom printed head bar were used to track head movements in relation to the target. Electrophysiology recordings were processed as described in the previous section. Matlab communicated with Cortex program online to control reward delivery every 500 ms if the mouse tracked the target for a duration greater than 1 s. The tracking threshold was defined as the center of the head within 50 mm in the x plane and 30 mm in the y plane of the target marker. Prior to recording sessions, mice were trained until they tracked the target consistently (∼1 hour of training over four days). Mice were first trained to track a target moving at a constant speed (16 mm/s). The speed of the target was then varied randomly during recording sessions (5-48 mm/s) and updated every 2 ms by Matlab. Self-rightward and -leftward movements were defined as movement to start the respective direction at least over 1 s. For linear regression analysis of single unit activity and behavioral variables, both neural data captured at 30 Hz and behavioral data captured at 100 Hz were split into 50 ms bins within a timeframe of 5 s in Neuroexplorer. Output data were exported into GraphPad Prism, where a linear regression analysis was performed. For population analyses, neural data and behavioral variables binned at 250 ms were standardized by a Z-score transformation within each session in Matlab. The transformed data from each animal were then averaged together to get a population Z-score for both the behavioral variables and the neural data. Once these Z-scores were obtained, the data were exported into GraphPad Prism, where a linear regression analysis was performed.

### Continuous Behavioral Classification

An unsupervised classification method was used to systematically and continuously measure behavior in pharmacological inactivation and PD optogenetic experiments. Animals were placed in an open-field arena and filmed with a camera positioned 0.7 meters from the floor of the arena. Data was recorded on a Dell computer (16GB RAM, Intel i7, 1TB SSD) via a 2.0 USB adaptor at 50 f/s. A region of interest was outlined and all further preprocessing was performed on the cropped image in Python using scientific tools and open source packages. Specifically, a binary threshold was applied to each frame of the video, and a 100 x 100 pixel frame was cropped around the animal. Each frame was then aligned (Berman, 2014 #47; https://github.com/gordonberman/MotionMapper) to eliminate allocentric components of behavior. Following the extraction and alignment of the mouse’s image, a wavelet decomposition was applied to increase the dimensionality of the image for better classification of postural dynamics. Next, a principal components analysis was performed to reduce the dimensionality to the top 10 principal components. The data was then fed into an autoregressive hidden Markov model (AR-HMM) ^21^. The AR-HMM model takes into account the image of the mouse, the duration of the specific behavioral ‘syllable,’ and the transition probabilities to other syllables. Specifically, using Gibbs sampling with an AR-HMM will segment all of the data into distinct behavioral states, and then switch to update these segments and the transitions between them ^40^. Each model goes through 1,000 iterations of Gibbs sampling. Importantly, all data (i.e. all groups) was co-trained to compare behavioral states across conditions. The output of the AR-HMM model allows behavioral classification into modules that occur systematically. Each video frame was assigned a behavioral label, which was used to calculate the probabilities of using each behavioral state, as well as the probabilities of transitioning between states. The model output was also used to compute forward and side velocities as previously described ^41^. Forward velocity is the component of a mouse’s velocity along the longitudinal axis of the body with respect to the position within previous video frame. Side velocity is the velocity component perpendicular to the longitudinal axis of the body with respect to the position within the previous video frame. Following the model output, additional statistical analyses were conducted. For each mouse, bootstrapped samples were computed to assess significance in module usage, as well as transition probabilities. Group comparisons were achieved with the Hotelling’s t-squared statistic. To assess individual differences between behavioral states, we estimated the variance of the data through bootstrapping. Using the bootstrapped estimates of variance, we performed a wald-test. We corrected for multiple comparisons using the Holm-Bonferonni method. Visualization of the data was also conducted in Python to create videos (OpenCV), graphs (networkx), and plots (matplotlib).

### Behavioral State and Transition Probability Analyses

Behavioral states were classified using an AR-HMM. Videos of the states were created and specific behaviors were characterized for the top 10 most commonly observed behaviors. Here the probabilities of displaying each behavior are sorted by degree of mobility. The probability of each behavior is displayed separately for each group. For the transition probability analysis, the probability of using certain behaviors is displayed in the size of the nodes. The probability of transitioning between behaviors is represented by the thickness of the lines. Positions of the states are kept constant across all three panels. Initial positions are seeded using a spring algorithm, where the probability of transferring between behaviors determines the spacing of the nodes. Nodes naturally repel each other. Each connection acts like a spring pulling together nodes with strength proportional to transition probability. The behavioral usage and transition probabilities were subtracted from each other, so that the remaining images represent the differences in probabilities between groups.

### Histology and Immunohistochemistry

Mice were deeply anesthetized and perfused with 0.1M PBS containing heparin followed by 4% paraformaldehyde after completion of experimentation. For mice with fiber and cannulae implants, heads were stored in 4% paraformaldehyde with 30% sucrose for 24-72 hrs at 4 °C to aid histological verification of placement. Brains were fixed in 30% sucrose thereafter. After sinking, brains were sliced coronally at 60 μm using a Leica CM1850 cryostat. For optogenetic experiments, the first 1-in-2 series of sections was processed for the presence of cytochrome oxidase to visualize cytoarchitecture. Briefly, sections were rinsed in 0.1M PB before incubating in a diaminobenzidine ^42^, cytochrome C, and sucrose solution for ∼2 hours at room temperature. The second series of sections were processed to enhance eYFP labeling. Sections were rinsed in 0.1M PBS for 20 min before being placed in a PBS-based blocking solution containing 5% goat serum and 0.25% Triton X-100 at room temperature for 1 hr. Sections were then incubated with a primary antibody (polyclonal chicken anti-GFP; 1:500 dilution; Abcam; catalog no. ab13970) in blocking solution overnight at 4 °C. Sections were then rinsed in PBS for 20 min before being placed in a secondary antibody used to visualize ChR2 as marked by YFP colocalization (goat anti-chicken Alexa Fluor 488; 1:1000 dilution; ThermoFisher; catalog no. A-11039) for 1 hr at room temperature. Fiber placement and injection site visualization was further aided by DAPI staining in the mounting medium (Fluoromount-G), or with the neuronal marker NeuN (monoclonal rabbit anti-NeuN; 1:1000 dilution; Abcam; catalog no. ab177487). For anatomical experiments, the endogenous signal caused by CAV2-Cre injection into *Ai14* mice was not enhanced. Primary antibodies for choline acetyltransferase (monoclonal rabbit anti-choline acetyltransferase; 1:1000 dilution; Abcam; catalog no. ab178850) and noradrenaline (polyclonal rabbit anti-noadrenaline; 1:500 dilution; Abcam; catalog no. ab8887) were used to identify neuronal subtype as marked by colocalization with tdTomato in the brainstem (goat anti-rabbit Alexa Fluor 488; 1:1000 dilution; Abcam; catalog no. ab150077) using the immunohistochemistry protocol previously described for eYFP enhancement. Cytochrome oxidase sections were dehydrated in 200 proof ethanol, defatted in xylene, and coverslipped with cytoseal. Sections for fluorescent microscopy were mounted and immediately coverslipped with Fluoromount aqueous mounting medium (Sigma; catalog no. F4680). Brightfield images were acquired and stitched using an Axio Imager.M1 upright microscope (Zeiss) and fluorescent images were acquired and stitched using a Z10 inverted microscope (Zeiss). Confirmation of optical fiber placement was performed by comparing images with a mouse brain atlas ^43^.

### Tyrosine Hydroxylase Processing and Quantification

A Tyrosine hydroxylase reaction was performed to visualize pallidal and striatal dopaminergic cell death in 6-OHDA injected mice. Briefly, sections were first agitated in a 3% H_2_O_2_ solution for 10 min at room temperature to block endogenous peroxidases. Sections were then transferred to a PBS-based blocking solution containing 5% goat serum and 0.25% Triton X-100 at room temperature for 1 hr. After incubating overnight in blocking solution containing a primary antibody (polyclonal rabbit anti-tyrosine hydroxylase; 1:1000 dilution; Millipore; catalog no. 657012), sections were incubated for 1 hr at room temperature in a biotinylated solution (biotinylated goat anti-rabbit IgG; 1:200 dilution; Vector Laboratories; catalog no. BA-1000) before incubating in an avidin-biotin horseradish peroxidase solution for 2 hrs at room temperature (Vector Novoxastra Laboratories). After rinsing sections in PBS, Tyrosine hydroxylase neurons were visualized with 0.05% DAB and 0.005% H_2_O_2_ in double distilled H_2_O, pH 7.2, for 5-10 min. The DAB reaction was stopped with subsequent PBS washes and stored at 4 °C. Tyrosine hydroxylase-reacted sections were dehydrated in 200 proof ethanol, defatted in xylene, and coverslipped with cytoseal. The intensity of tyrosine hydroxylase staining in saline injected (n = 3) and bilateral-PD experimental groups (n = 10) was quantified as previously described ^20^. Bilateral-PD animals were included in analyses if tyrosine hydroxylase intensity values were <80% compared to controls. Two bilateral-PD animals were excluded from the study because they did not meet this criterion.

### Quantification and Statistical Analysis

The values and error bars reported in the text are the mean ± SEM, respectively. Behavioral data was analyzed with MATLAB 2016b (MathWorks). Statistical tests were performed in Prism 8 (GraphPad). Two-tailed parametric tests were used. An *a priori* alpha level of 0.05 was used to determine significance.

## Supplementary Figures

**Supplemental Figure 1.**
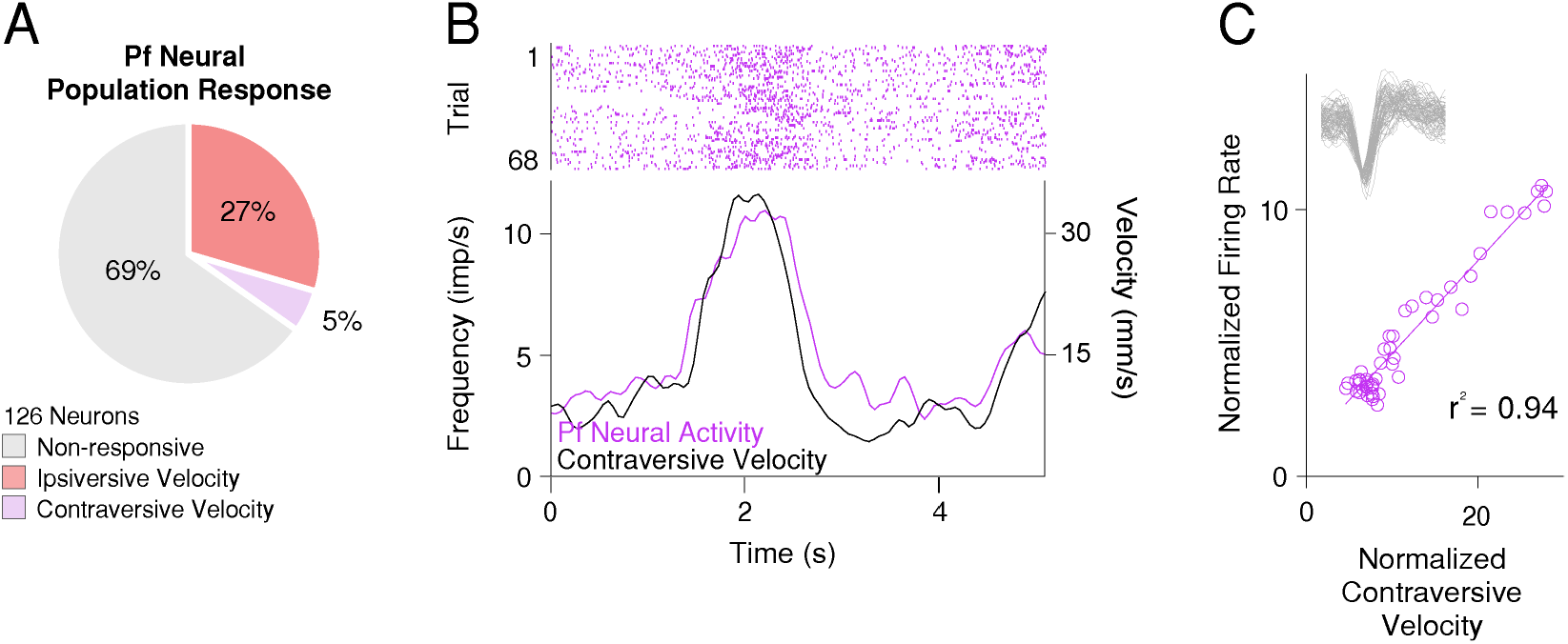
A Subset of Pf Neurons Represent Contraversive Velocity. **(A)** Pf neural population response during reward tracking (*n* = 126). The majority of neurons in the Pf did not correlate with movement kinematics (69%). The remaining neurons represent ipsiversive (27%) and contraversive (5%) velocity. **(B)** Peri-event histogram showing representative example of contraversive encoding neuron in Pf (purple raster and trace, left panel). **(C)** Linear regression analysis (*r*^2^ *=* 0.94) of Pf firing frequency and contraversive velocity of representative neuron (right panel).

**Supplemental Figure 2.**
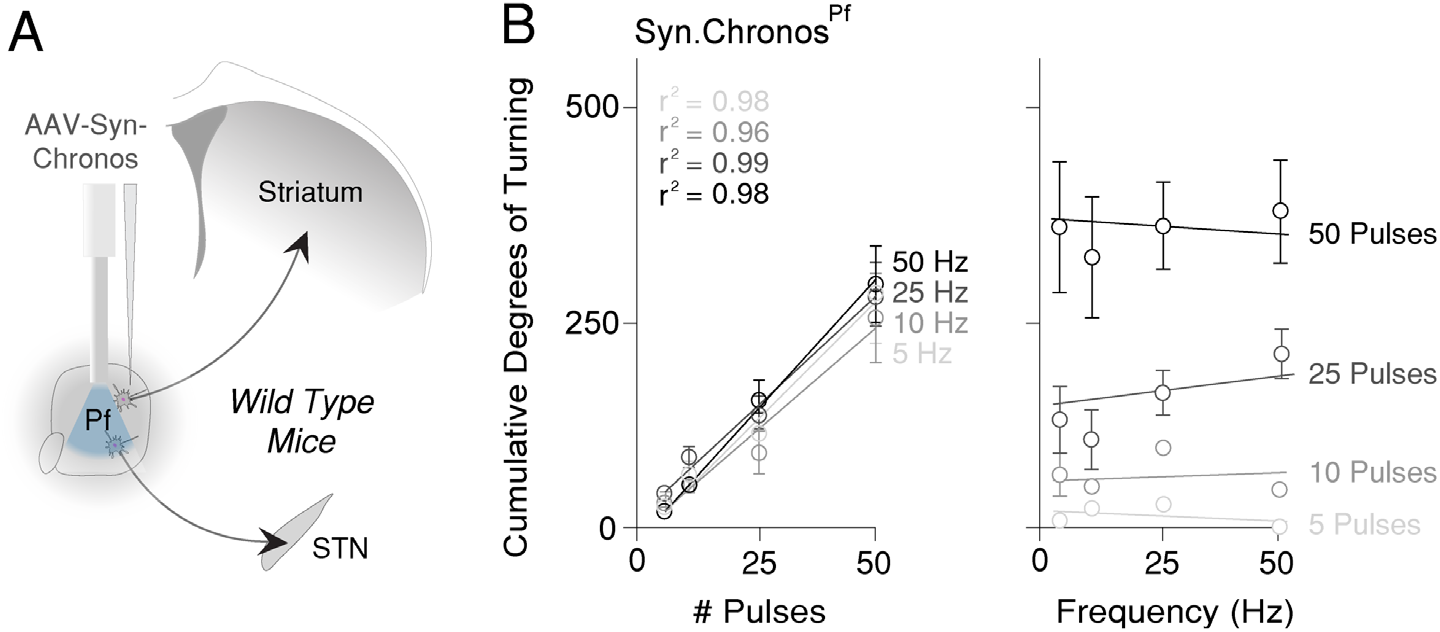
Pf Cell-Type Nonspecific Excitation Causes Ipsiversive Turning across Stimulation Frequencies. **(A)** Optogenetic excitation of neurons in the Pf (AAV5.Syn.Chronos-GFP.WPRE.bGH, *wild type* mice, *n =* 7). **(B)** Cumulative rotation in degrees significantly increase as a function of the number of excitation pulses (left), regardless of the excitation frequency (right) (Linear regression analyses, all *r*^2^ ≥ 0.96). Error bars = mean ± SEM.

**Supplemental Figure 3.**
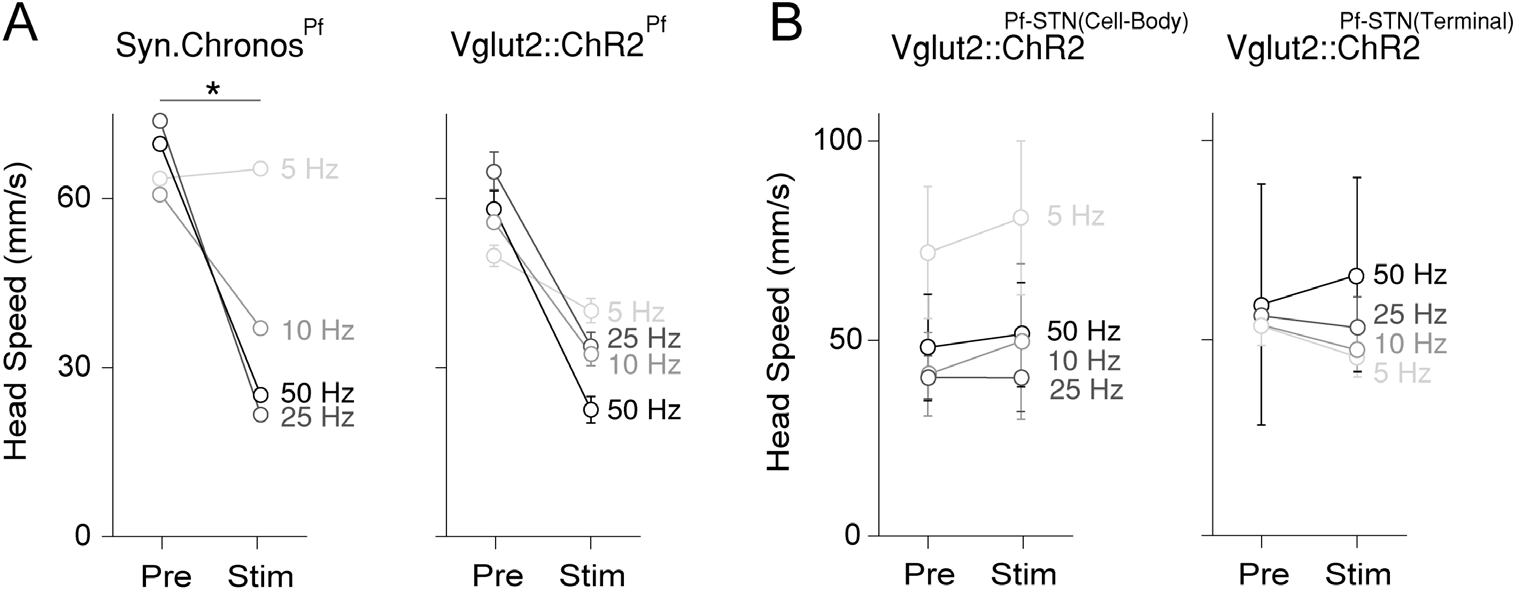
Bilateral Pf Cell-Body, but not Pf-STN Pathway Specific Optogenetic Excitation Slows Movement. (**A**) Head speed during bilateral optogenetic stimulation of Pf neurons across various frequencies. Head speed is significantly decreased due to excitation in Syn.Chronos animals (two-way RM ANOVA (*F*(1,2) = 58.84, *p* = 0.0166, *n* = 3)). A difference in head speed approached significance in Vglut2::ChR2 animals (two-way RM ANOVA, pre vs. stimulation (*F*(1,2) = 7.721, *p* = 0.1088, *n* = 3)). (**B**) Bilateral terminal excitation (two-way RM ANOVA (*F*(1,2) = 1.011, *p* = 0.4206, *n* = 3)) and cell-body excitation (two-way RM ANOVA *F*(1,2) = 5.434, *p* = 0.1450, *n* = 3)) of Pf-STN neurons did not significantly influence head speed. Error bars = mean ± SEM.

**Supplemental Figure 4.**
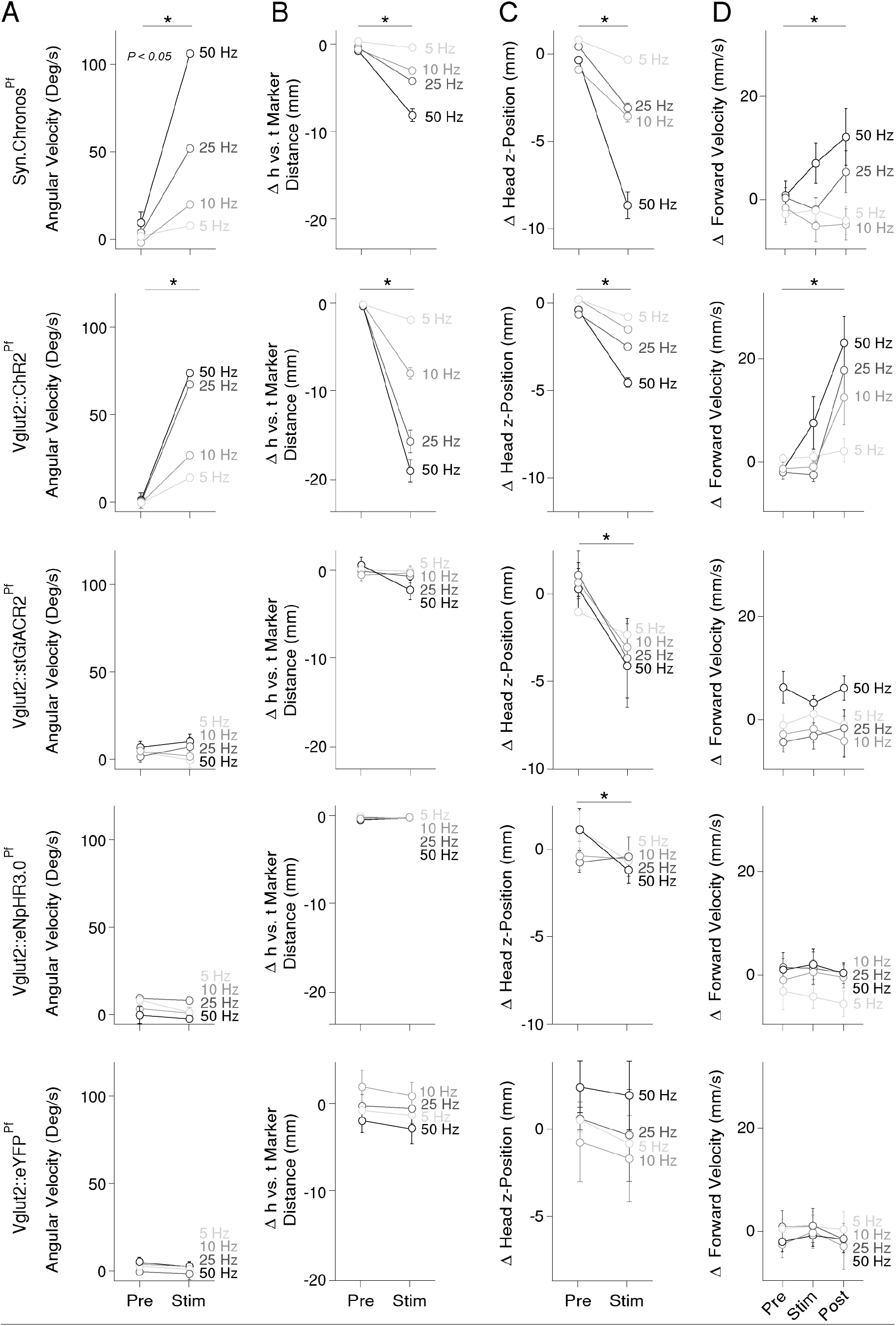
Pf Unilateral Optogenetic Excitation, but not Inhibition Causes Ipsiversive Turning and Postural Changes across Stimulation Frequencies. **(A)** Unilateral optogenetic excitation of Pf significantly increased angular velocity in Syn.Chronos (two-way RM ANOVA, pre vs. stimulation (*F*(1,6) = 16.68, *p* = 0.0065, *n* = 7)) and Vglut2::ChR2 (two-way RM ANOVA, pre vs. stimulation (*F*(1,5) = 12.46, *p* = 0.0167, *n* = 6)) animals across various frequencies. There was no significant decrease in Vglut2::stGtACR2 (two-way RM ANOVA, pre vs. stimulation (*F*(1,5) = 0.19808, *p* = 0.6804, *n* = 6)), Vglut2::eNpHR3.0 (two-way RM ANOVA, pre vs.stimulation (*F*(1,5) = 1.84, *p* = 0.2330, *n* = 6)) and Vglut2::eYFP (two-way RM ANOVA, pre vs. stimulation (*F*(1,5) = 17.36, *p* = 0.0088, Sidak’s *post hoc* multiple comparison test revealed no significant effect of frequency pre vs. post stimulation (*p* > 0.20), *n* = 6)) animals **(B)** Unilateral optogenetic excitation of Pf caused a significant bending of the torso in Syn.Chronos (two-way RM ANOVA, pre vs. stimulation (*F*(1,6) = 16.62, *p* = 0.0065, *n* = 7)) and Vglut2::ChR2 (two-way RM ANOVA, pre vs. stimulation (*F*(1,5) = 21.59, *p* = 0.0056, *n* = 6)) animals. There was no significant decrease in Vglut2::stGtACR2 (two-way RM ANOVA, pre vs. stimulation (*F*(1,5) = 2.772, *p* = 0.1568, *n* = 6)), Vglut2::eNpHR3.0 (two-way RM ANOVA, pre vs. stimulation (*F*(1,5) = 0.02698, *p* = 0.8760, *n* = 6)and Vglut2::eYFP (two-way RM ANOVA, pre vs. stimulation (*F*(1,5) = 3.603, *p* = 0.1161, *n* = 6)) animals **(C)** Unilateral optogenetic excitation of Pf caused a significant head elevation change in Syn.Chronos (two-way RM ANOVA, pre vs. stimulation (*F(*1,6) = 9.617, *p* = 0.0211, *n* = 7)), Vglut2::stGtACR2 (two-way RM ANOVA, pre vs. stimulation (*F*(1,5) = 13.47, *p* = 0.0144, *n* = 6)), and Vglut2::eNpHR3.0 (two-way RM ANOVA, pre vs. stimulation (*F*(1,5) = 11.15, *p* = 0.0206, *n* = 6) animals. There was no significant change in Vglut2::ChR2 animals (two-way RM ANOVA, pre vs. stimulation (*F*(1,5) = 21.59, *p* = 0.0056, *n* = 6)), however Sidak’s *post hoc* multiple comparison test revealed a significant difference between pre-stimulation and the 50 Hz condition (*t*(15) = 4.69, *p* = 0.0010)). There was also no significant decrease in Vglut2::eYFP (two-way RM ANOVA, pre vs. stimulation (*F*(1,5) = 1.265, *p* = 0.3118, *n* = 6)) animals. **(D)** Unilateral optogenetic excitation caused a significant change in forward velocity in both Syn.Chronos (two-way RM ANOVA, epoch (*F(*2,12) = 3.59, *p* = 0.06, *n* = 7)) and Vglut2 (two-way RM ANOVA, epoch (*F(*2,10) = 16.69, *p* = 0.0007, *n* = 6)) animals post-stimulation. There was no significant increase in Vglut2::stGtACR2 (two-way RM ANOVA, epoch (*F(*2,10) = 0.05138, *p* = 0.9502, *n* = 6)), Vglut2::eNpHR3.0 (two-way RM ANOVA, epoch (*F(*2,10) = 2.362, *p* = 0.1445, *n* = 6)), and Vglut2::eYFP (two-way RM ANOVA, epoch (*F(*2,10) = 5.139, *p* = 0.0292, Sidak’s *post hoc* multiple comparison test revealed no significant between epochs across stimulation frequencies (*p* > 0.20), *n* = 6)). Error bars = mean ± SEM.

**Supplemental Figure 5.**
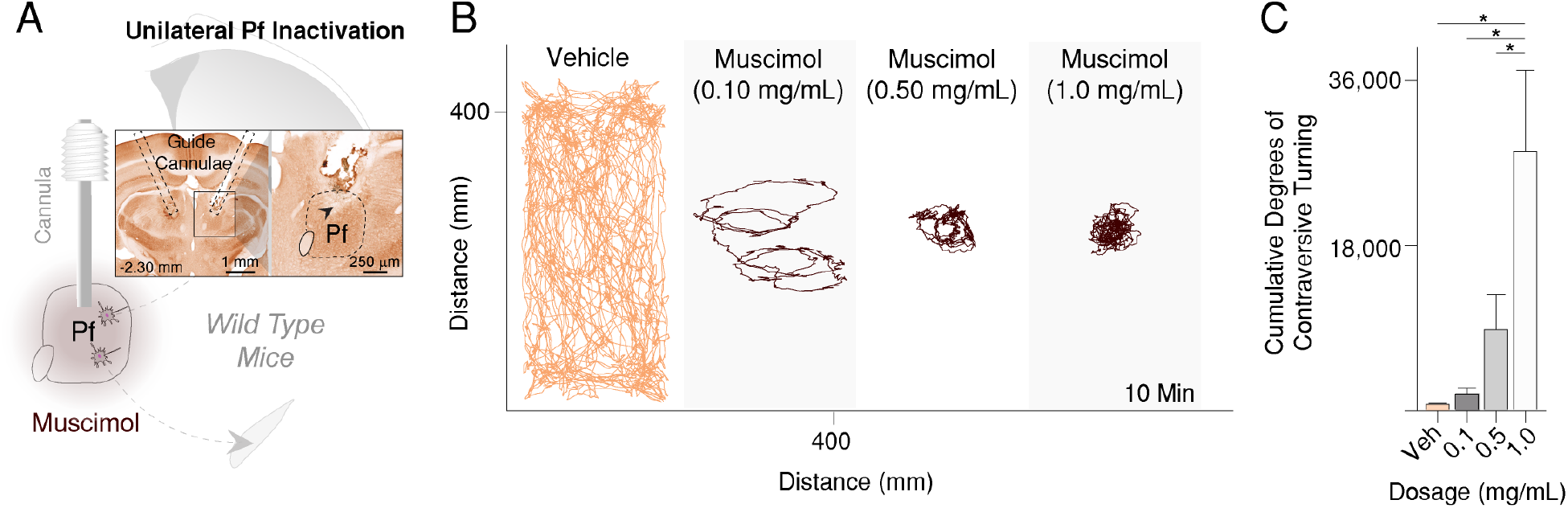
Unilateral Pf Pharmacological Inhibition Produces Contraversive Turning. (**A**) Pf unilateral muscimol inactivation through chronically implanted guide cannulae. Coronal section through the thalamus shows cannula placement above the Pf bilaterally. Black arrow indicates guide cannula tip. (**B**) Representative example of movement traces after unilateral vehicle and muscimol (0.10, 0.50, and 1.0 µg/µL) injections into the Pf. (**C**) Cumulative rotations in degrees significantly increased at higher muscimol dosages (one-way RM ANOVA, significant main effect of dosage on cumulative rotations across conditions, *F*(3,28) = 6.944, *p* = 0.0012, *n* = 8). *Post hoc* comparison using Tukey’s multiple comparison test indicated that the mean cumulative rotations for the 1.0 µg/µL dosage condition (M = 78.5, SD = 69.7) was significantly different from the vehicle (M = 2.0, SD = 0.93), 0.10 µg/µL (M = 5.1, SD = 5.3), and 0.50 µg/µL (M = 24.6, SD = 29.9) conditions. The 0.10 µg/µL and 0.50 µg/µL conditions were not significantly different from the vehicle condition.

**Supplemental Figure 6.**
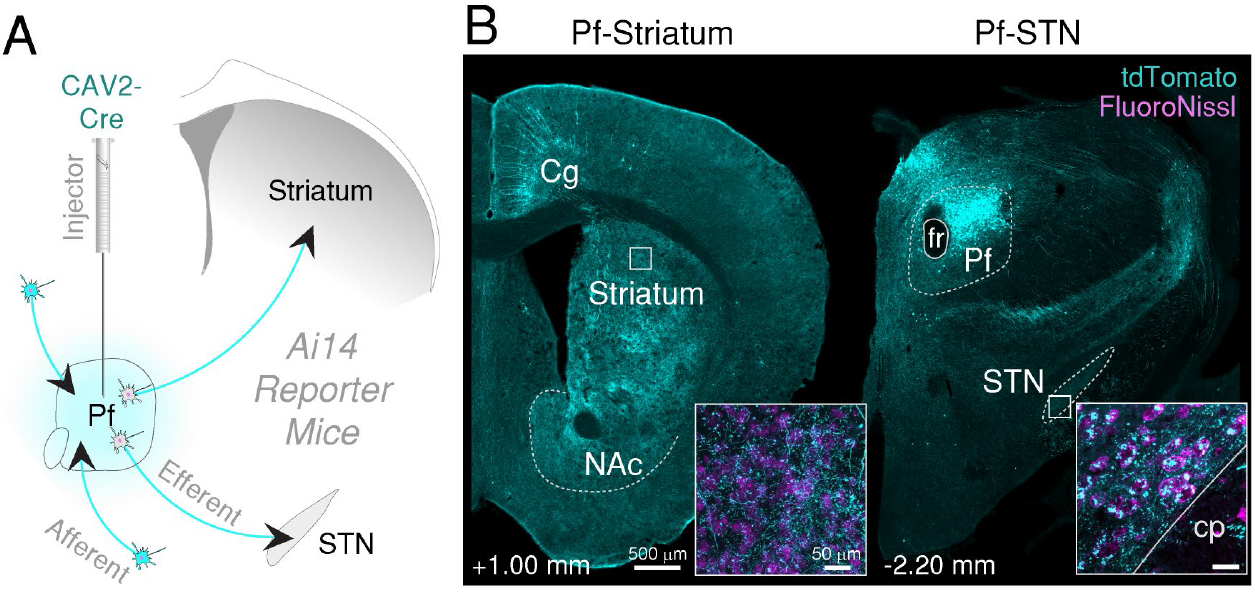
Pf Projects to the Striatum and STN in Mice. (**A**) CAV2-Cre injection into the Pf of *Ai14* transgenic mice to visualize afferent and efferent projections. A high-titer injection of CAV2-Cre allows visualization of input-output connectivity. Neurons in *Ai14* mice fluoresce (tdTomato, cyan) when infected with a virus with Cre-recombinase. Pf injection site shown in Figure 7. (**B**) Pf terminal labeling in the striatum and STN. Insets show high magnification view of Pf terminal labeling. Abbreviations – Cg, cingulate cortex; cp, cerebral peduncle; fr, fasciculus retroflexus; NAc, nucleus accumbens.

**Supplemental Figure 7.**
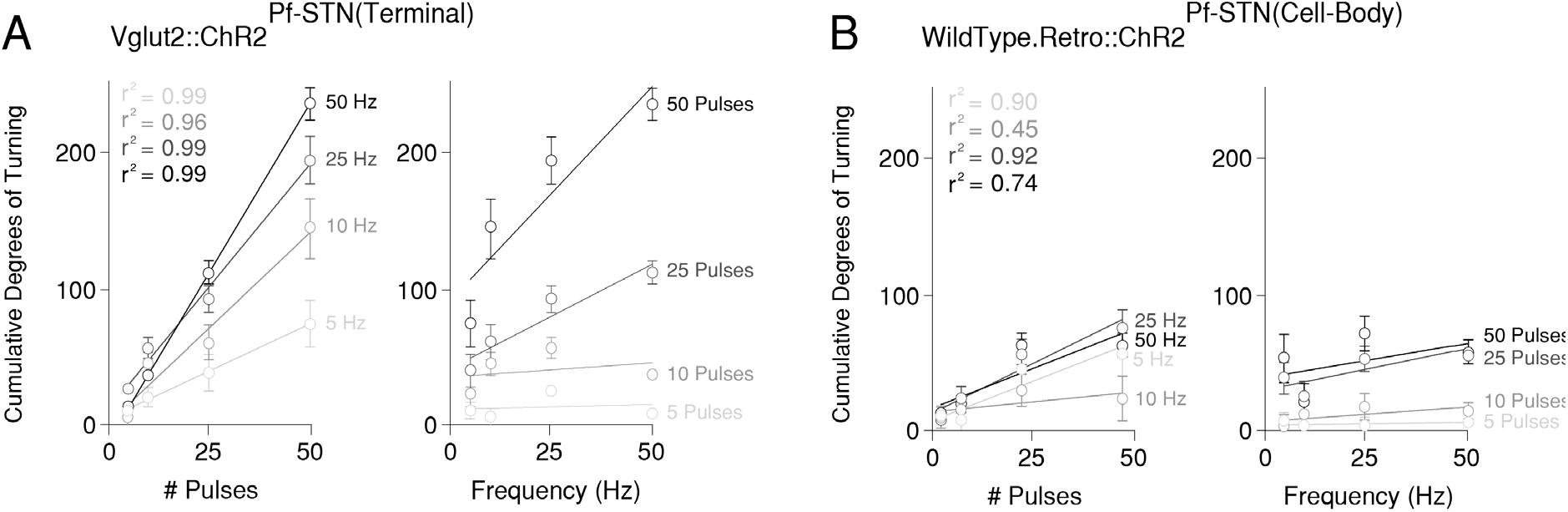
Pf-STN Optogenetic Excitation Produces Ipsiversive Turning. **(A)** Optogenetic terminal excitation of *Vglut2+* Pf-STN neurons. Cumulative rotation in degrees significantly increased as a function of the number of excitation pulses (left, linear regression analyses, all *r*^2^ ≥ 0.96). **(B)** Optogenetic cell-body excitation of Pf-STN neurons in *wild type* mice. Cumulative rotation in degrees significantly increased at 25 Hz and 50 Hz as a function of the number of excitation pulses (left, linear regression analyses, all *r*^2^ ≥ 0.90). Error bars = mean ± SEM.

**Supplemental Figure 8.**
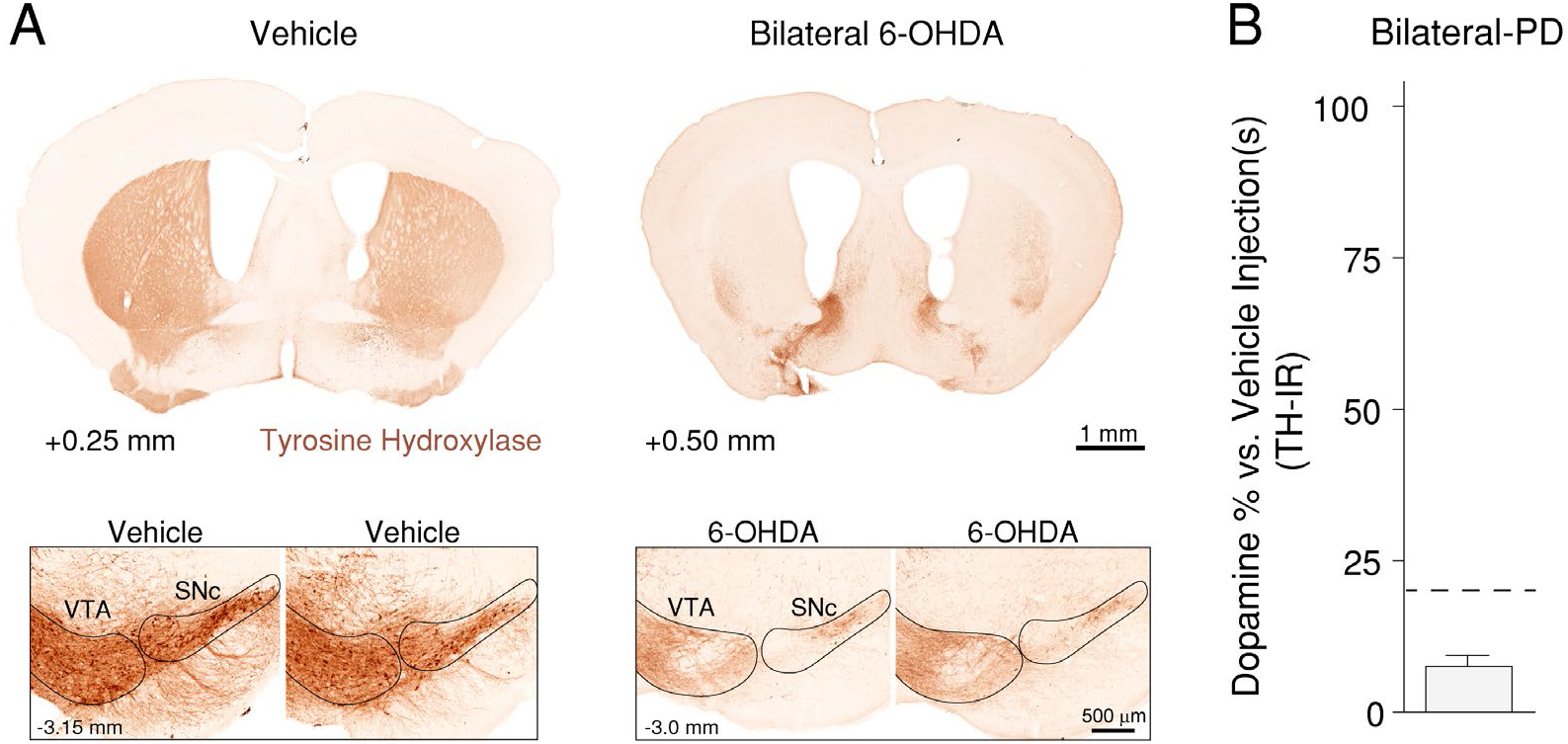
Parkinsonian Mouse Model Verification through Tyrosine Hydroxylase Quantification. **(A)** Representative coronal sections showing TH immunoreactivity in the striatum of control (left) and bilateral-PD (right) mice. Bottom insets show TH reactivity in the SNc for each group. **(B)** TH immunoreactivity quantification in bilateral-PD experiments. Animals were considered Parkinsonian if TH immunoreactivity was >80% compared to vehicle injected control animals. Abbreviations – SNc, substantia nigra pars compacta; VTA, ventral tegmental area. Error bars = mean ± SEM.

**Supplemental Figure 9.**
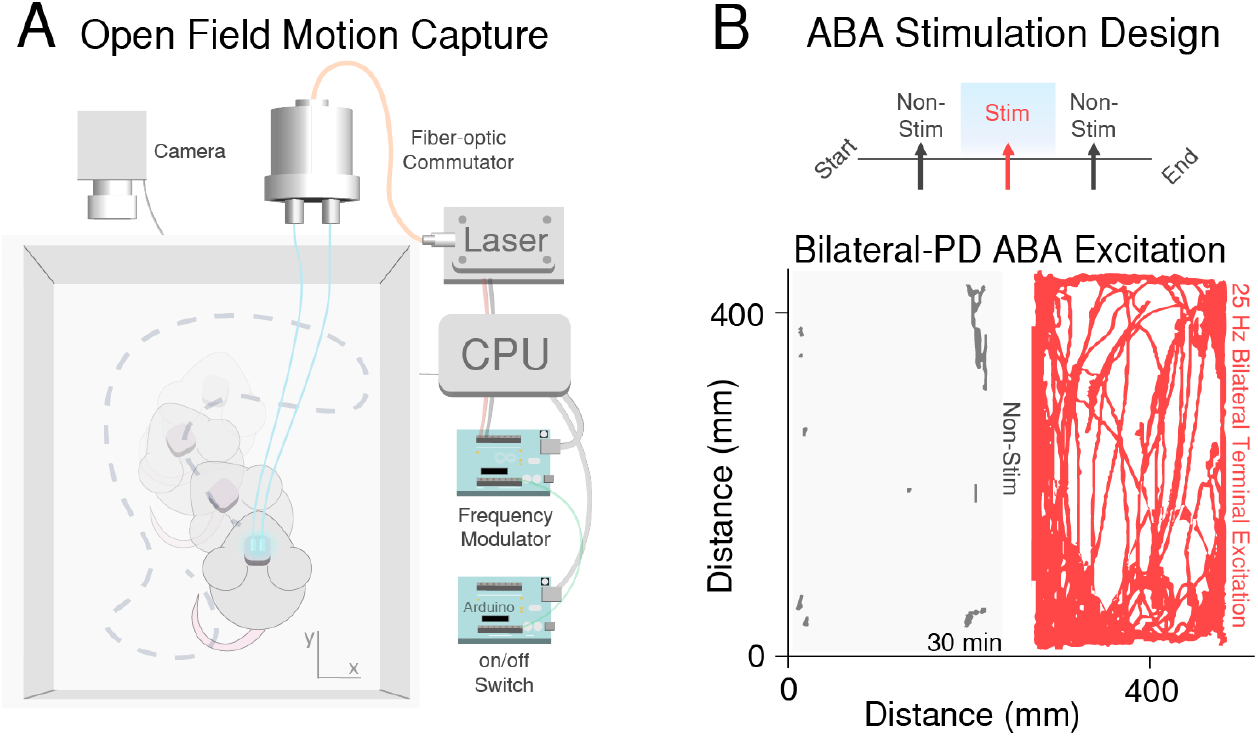
Experimental Design for Parkinsonian Rescue Experiments. **(A)** 2D open field video capture of Parkinsonian mice during optogenetic stimulation. Two Arduinos controlled laser output through optic patch cables connected to implanted optic fibers over the Pf or STN. **(B)** ABA stimulation design (top). Optogenetic stimulation epochs (red) were interspersed between non-stimulation epochs (grey). For bilateral-PD experiments, the ABA stimulation session lasted for an hour (1 min on, 1 min off). Representative open-field movement traces from bilateral Pf-STN terminal optogenetic excitation experiment (30 x 1 min stimulation and non-stimulation epochs). Optogenetic excitation increased locomotion during stimulation epochs (red) compared to non-stimulation epochs (grey).

**Supplemental Figure 10.**
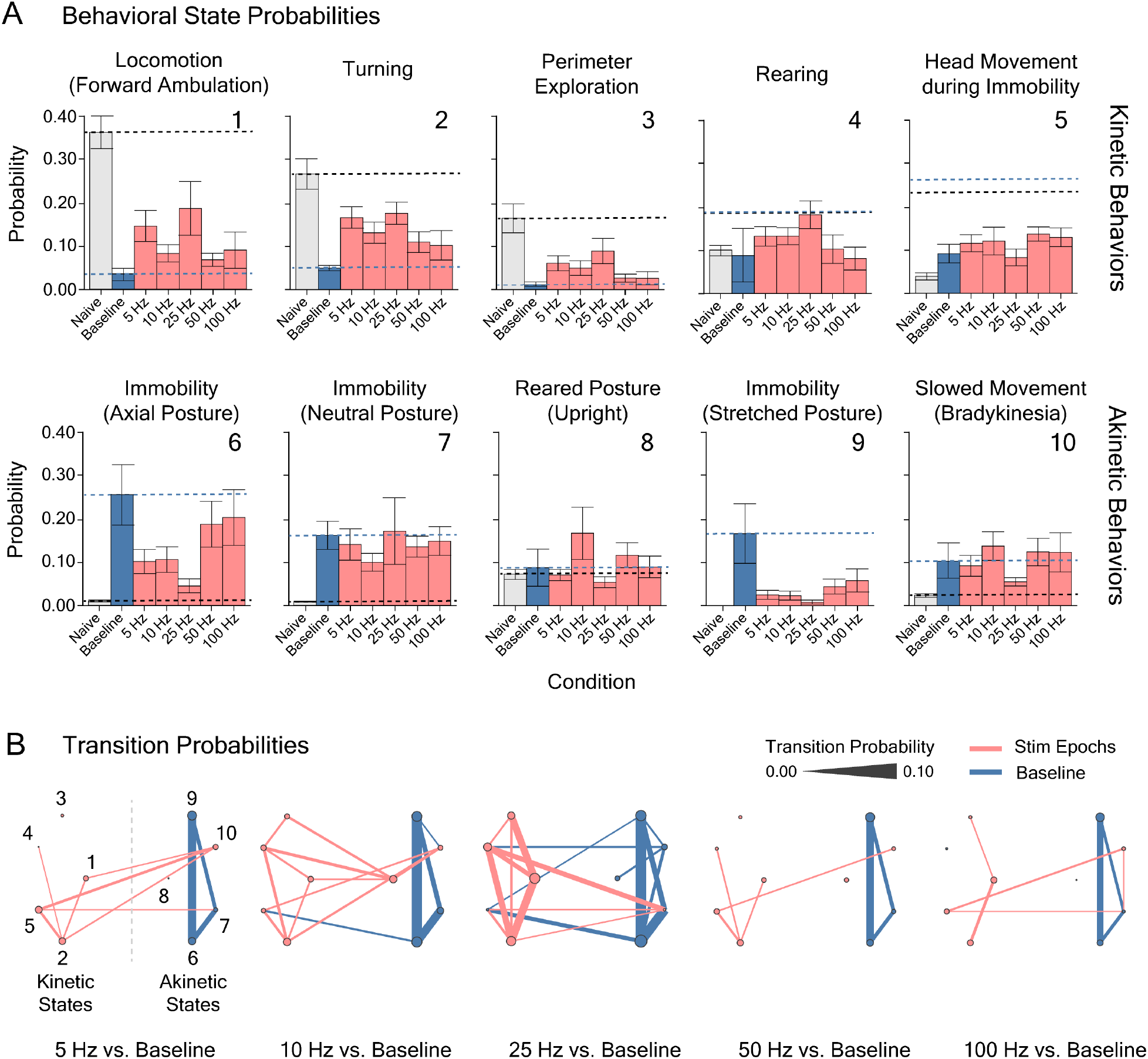
Optogenetic Excitation of Pf-STN Cell-Bodies in Bilateral 6-OHDA Parkinsonian Mice Restores Natural Behaviors as Revealed by an AR-HMM. (**A**) Probability of usage of AR-HMM identified kinetic (top row) and akinetic (bottom row) behavioral states during Pf-STN cell-body excitation epochs (ChR2, red) across various frequencies (5, 10, 25, 50, and 100 Hz) in bilateral 6-OHDA PD mice. Stimulation groups are statistically different from the akinetic baseline condition (blue) (Hotelling’s t-tests, *p* < 0.0001 for all comparisons except 5 Hz (*p* > 0.05)). Error bars = SEM. Dotted lines represent mean in akinetic baseline and naïve conditions. Numbers represent behavioral states shown in (C). (**B**) Pf-STN cell-body excitation increases mobility between kinetic behavioral states compared to the akinetic baseline in the 25 Hz condition in bilateral-PD mice. Bigrams illustrating transition probabilities between AR-HMM identified kinetic (left) and akinetic (right) states across frequencies during Pf-STN cell-body excitation epochs (red) versus the akinetic baseline condition (blue). Line thickness represents transition probability. Numbers represent behavioral states identified in (B): 1) locomotion (forward ambulation); 2) turning; 3) perimeter exploration; 4) rearing; 5) head movement during immobility; 6) immobility (axial posture); 7) immobility (neutral posture); 8) reared posture; 9) immobility (stretched posture); 9) slowed movement (bradykinesia). Error bars = mean ± SEM.

**Supplemental Figure 11.**
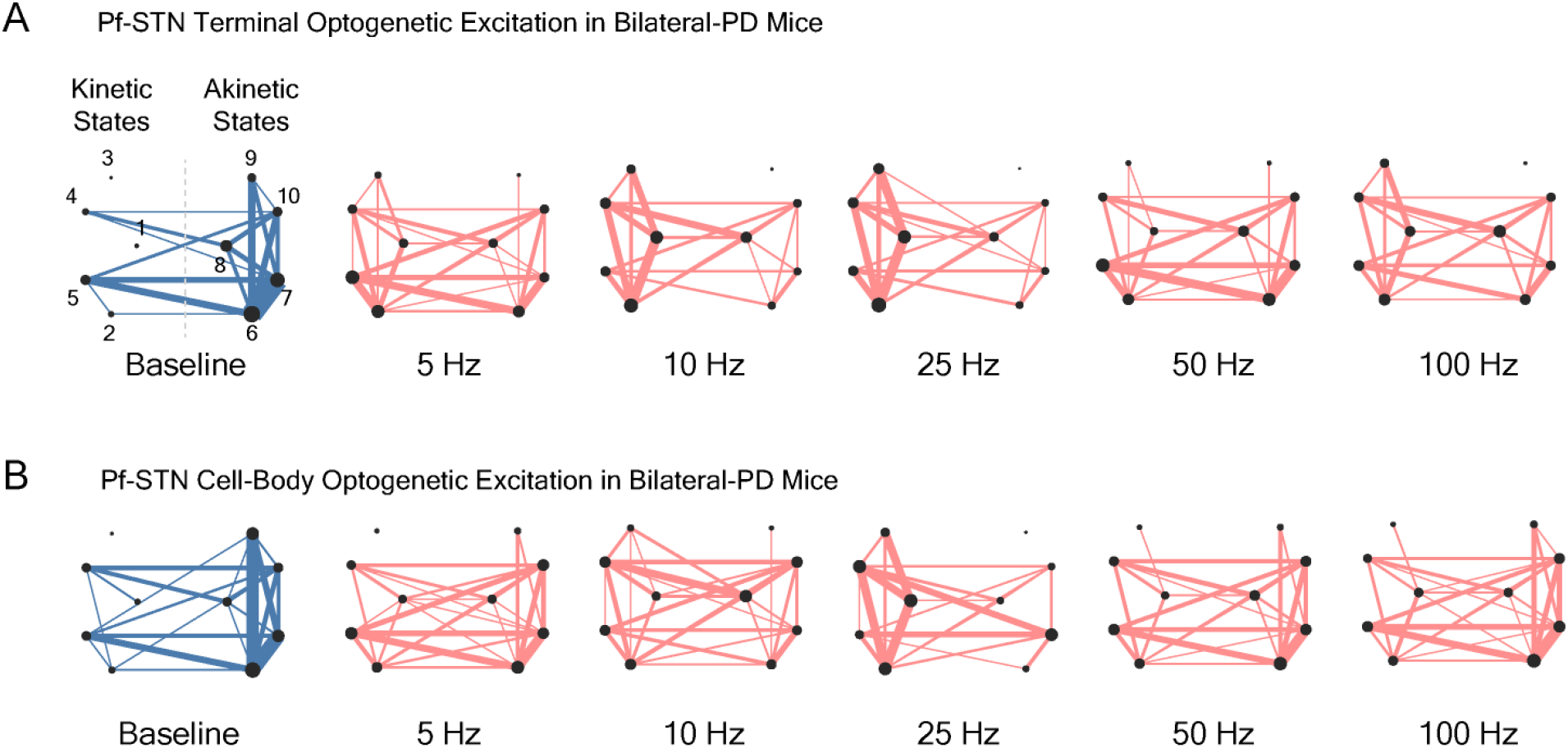
Bigrams used in transition probability analysis before subtraction. **(A)** Transition probability bigrams from Pf-STN terminal optogenetic excitation experiments (Figure 5). Baseline bigram (blue, left) presented with stimulation bigrams arranged by frequency (red, right). Bigrams spatially separated based on kinetic (left, 1-5) and akinetic (right, 6-10) identified by an AR-HMM. See Figure 5 for corresponding behavioral states. **(B)** Transition probability bigrams from Pf-STN cell-body optogenetic excitation experiments (Figure S12). Baseline bigram (blue, left) presented with stimulation bigrams arranged by frequency (red, right). Bigrams spatially separated based on kinetic (left, 1-5) and akinetic (right, 6-10) identified by an AR-HMM. See Figure 5 for corresponding behavioral states.

**Supplemental Figure 12.**
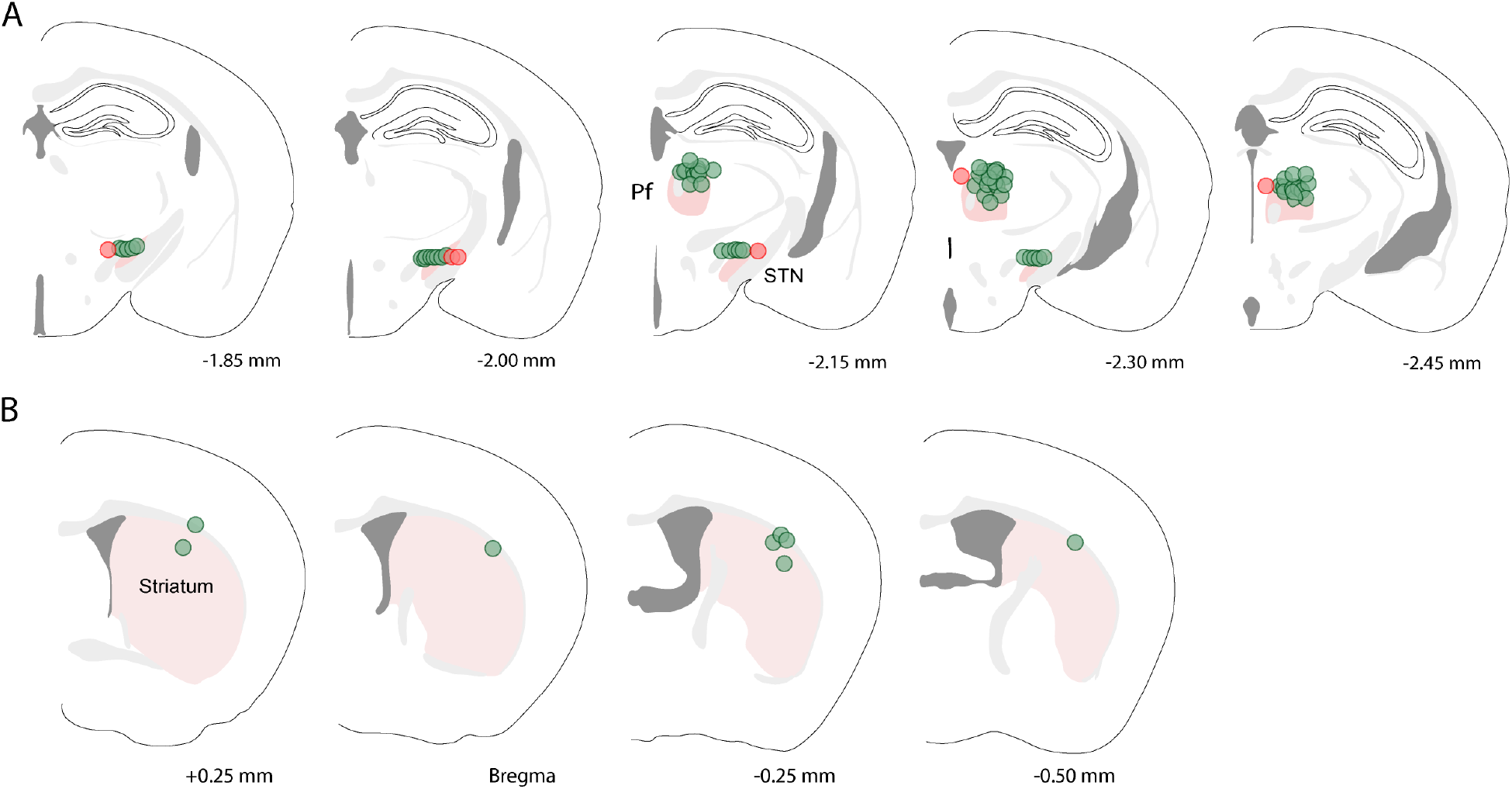
Optic fiber placement shown in coronal sections. **(A)** Coronal sections through the diencephalon showing optic fiber placements into the Pf and STN. Green circles denote included data points. Red circles denote excluded data points. **(B)** Coronal section through the striatum showing optic fiber placements into the dorsal striatum. Bregma indicated below sections.

## Supplementary Movies

Movie S1. Pf *Vglut2+* unilateral optogenetic excitation

Movie S2. Pf unilateral muscimol inactivation

Movie S3. Unilateral Pf-STN *Vglut2+* terminal optogenetic excitation

Movie S4. Bilateral Pf-STN *Vglut2+* terminal optogenetic excitation in bilateral-PD

Movie S5. Bilateral Pf-STN cell-body optogenetic excitation in bilateral-PD

Movie S6. Auto-Regressive Hidden Markov behavioral state classification for PD experiments

## References

1. Wilson, C. R. G. C. J. Chapter II The basal Ganglia. Vol. 12 371–468 (Elsevier, 1996).

2. Diaz-Hernandez, E. et al. The Thalamostriatal Projections Contribute to the Initiation and Execution of a Sequence of Movements. Neuron 100, 739–752.e735, doi:10.1016/j.neuron.2018.09.052 (2018).

3. Ding, J. B., Guzman, J. N., Peterson, J. D., Goldberg, J. A. & Surmeier, D. J. Thalamic gating of corticostriatal signaling by cholinergic interneurons. Neuron 67, 294–307, doi:10.1016/j.neuron.2010.06.017 (2010).

4. Parker, P. R., Lalive, A. L. & Kreitzer, A. C. Pathway-Specific Remodeling of Thalamostriatal Synapses in Parkinsonian Mice. Neuron 89, 734–740, doi:10.1016/j.neuron.2015.12.038 (2016).

5. Matsumoto, N., Minamimoto, T., Graybiel, A. M. & Kimura, M. Neurons in the thalamic CM-Pf complex supply striatal neurons with information about behaviorally significant sensory events. Journal of neurophysiology 85, 960–976, doi:10.1152/jn.2001.85.2.960 (2001).

6. Brown, H. D., Baker, P. M. & Ragozzino, M. E. The parafascicular thalamic nucleus concomitantly influences behavioral flexibility and dorsomedial striatal acetylcholine output in rats. The Journal of neuroscience: the official journal of the Society for Neuroscience 30, 14390–14398, doi:10.1523/jneurosci.2167-10.2010 (2010).

7. Tanimura, A., Du, Y., Kondapalli, J., Wokosin, D. L. & Surmeier, D. J. Cholinergic Interneurons Amplify Thalamostriatal Excitation of Striatal Indirect Pathway Neurons in Parkinson’s Disease Models. Neuron, doi:10.1016/j.neuron.2018.12.004 (2019).

8. Smith, Y. et al. The thalamostriatal system in normal and diseased states. Frontiers in systems neuroscience 8, 5, doi:10.3389/fnsys.2014.00005 (2014).

9. Kita, T., Shigematsu, N. & Kita, H. Intralaminar and tectal projections to the subthalamus in the rat. The European journal of neuroscience 44, 2899–2908, doi:10.1111/ejn.13413 (2016).

10. Fan, D. et al. A wireless multi-channel recording system for freely behaving mice and rats. PloS one 6, e22033, doi:10.1371/journal.pone.0022033 (2011).

11. Bartholomew, R. A. et al. Striatonigral control of movement velocity in mice. The European journal of neuroscience 43, 1097–1110, doi:10.1111/ejn.13187 (2016).

12. Kim, N., Barter, J. W., Sukharnikova, T. & Yin, H. H. Striatal firing rate reflects head movement velocity. The European journal of neuroscience 40, 3481–3490, doi:10.1111/ejn.12722 (2014).

13. Fremeau, R. T., Jr. et al. The expression of vesicular glutamate transporters defines two classes of excitatory synapse. Neuron 31, 247–260 (2001).

14. Lei, W. et al. Confocal laser scanning microscopy and ultrastructural study of VGLUT2 thalamic input to striatal projection neurons in rats. The Journal of comparative neurology 521, 1354–1377, doi:10.1002/cne.23235 (2013).

15. Wiegert, J. S., Mahn, M., Prigge, M., Printz, Y. & Yizhar, O. Silencing Neurons: Tools, Applications, and Experimental Constraints. Neuron 95, 504–529, doi:10.1016/j.neuron.2017.06.050 (2017).

16. Mahn, M. et al. High-efficiency optogenetic silencing with soma-targeted anion-conducting channelrhodopsins. Nature communications 9, 4125, doi:10.1038/s41467-018-06511-8 (2018).

17. Di Chiara, G., Morelli, M., Porceddu, M. L. & Gessa, G. L. Role of thalamic gamma-aminobutyrate in motor functions: catalepsy and ipsiversive turning after intrathalamic muscimol. Neuroscience 4, 1453–1465 (1979).

18. Tervo, D. G. et al. A Designer AAV Variant Permits Efficient Retrograde Access to Projection Neurons. Neuron 92, 372–382, doi:10.1016/j.neuron.2016.09.021 (2016).

19. Thiele, S. L., Warre, R. & Nash, J. E. Development of a unilaterally-lesioned 6-OHDA mouse model of Parkinson’s disease. Journal of visualized experiments: JoVE, doi:10.3791/3234 (2012).

20. Mastro, K. J. et al. Cell-specific pallidal intervention induces long-lasting motor recovery in dopamine-depleted mice. Nature neuroscience 20, 815–823, doi:10.1038/nn.4559 (2017).

21. Wiltschko, A. B. et al. Mapping Sub-Second Structure in Mouse Behavior. Neuron 88, 1121–1135, doi:10.1016/j.neuron.2015.11.031 (2015).

22. Zingg, B. et al. AAV-Mediated Anterograde Transsynaptic Tagging: Mapping Corticocollicular Input-Defined Neural Pathways for Defense Behaviors. Neuron 93, 33–47, doi:10.1016/j.neuron.2016.11.045 (2017).

23. Porceddu, M. L., Piacente, B., Morelli, M. & Di Chiara, G. Opposite turning effects of dainic and ibotenic acid injected in the rat substantia nigra. Neuroscience letters 15, 271–276 (1979).

24. Barter, J. W., Castro, S., Sukharnikova, T., Rossi, M. A. & Yin, H. H. The role of the substantia nigra in posture control. The European journal of neuroscience 39, 1465–1473, doi:10.1111/ejn.12540 (2014).

25. Bradfield, L. A. & Balleine, B. W. Thalamic Control of Dorsomedial Striatum Regulates Internal State to Guide Goal-Directed Action Selection. The Journal of neuroscience: the official journal of the Society for Neuroscience 37, 3721–3733, doi:10.1523/jneurosci.3860-16.2017 (2017).

26. Hunnicutt, B. J. et al. A comprehensive excitatory input map of the striatum reveals novel functional organization. eLife 5, doi:10.7554/eLife.19103 (2016).

27. Kravitz, A. V. et al. Regulation of parkinsonian motor behaviours by optogenetic control of basal ganglia circuitry. Nature 466, 622–626, doi:10.1038/nature09159 (2010).

28. Gradinaru, V., Mogri, M., Thompson, K. R., Henderson, J. M. & Deisseroth, K. Optical deconstruction of parkinsonian neural circuitry. Science (New York, N.Y.) 324, 354–359, doi:10.1126/science.1167093 (2009).

29. Mastro, K. J., Bouchard, R. S., Holt, H. A. & Gittis, A. H. Transgenic mouse lines subdivide external segment of the globus pallidus (GPe) neurons and reveal distinct GPe output pathways. The Journal of neuroscience: the official journal of the Society for Neuroscience 34, 2087–2099, doi:10.1523/jneurosci.4646-13.2014 (2014).

30. Schaafsma, J. D. et al. Characterization of freezing of gait subtypes and the response of each to levodopa in Parkinson’s disease. European journal of neurology 10, 391–398 (2003).

31. Schaafsma, J. D. et al. Gait dynamics in Parkinson’s disease: relationship to Parkinsonian features, falls and response to levodopa. Journal of the neurological sciences 212, 47–53 (2003).

32. Nonnekes, J. et al. Unmasking levodopa resistance in Parkinson’s disease. Movement disorders: official journal of the Movement Disorder Society 31, 1602–1609, doi:10.1002/mds.26712 (2016).

33. Remple, M. S. et al. Subthalamic nucleus neuronal firing rate increases with Parkinson’s disease progression. Movement disorders: official journal of the Movement Disorder Society 26, 1657–1662, doi:10.1002/mds.23708 (2011).

34. Miocinovic, S., Somayajula, S., Chitnis, S. & Vitek, J. L. History, applications, and mechanisms of deep brain stimulation. JAMA neurology 70, 163–171, doi:10.1001/2013.jamaneurol.45 (2013).

35. Mazzone, P. et al. Bilateral Implantation of Centromedian-Parafascicularis Complex and GPi: A New Combination of Unconventional Targets for Deep Brain Stimulation in Severe Parkinson Disease. Neuromodulation: journal of the International Neuromodulation Society 9, 221–228, doi:10.1111/j.1525-1403.2006.00063.x (2006).

36. Anderson, D., Beecher, G. & Ba, F. Deep Brain Stimulation in Parkinson’s Disease: New and Emerging Targets for Refractory Motor and Nonmotor Symptoms. Parkinson’s disease 2017, 5124328, doi:10.1155/2017/5124328 (2017).

37. Lanciego, J. L. et al. The search for a role of the caudal intralaminar nuclei in the pathophysiology of Parkinson’s disease. Brain research bulletin 78, 55–59, doi:10.1016/j.brainresbull.2008.08.008 (2009).

38. Nambu, A., Tokuno, H. & Takada, M. Functional significance of the cortico-subthalamo-pallidal ‘hyperdirect’ pathway. Neuroscience research 43, 111–117 (2002).

39. Lopes, G. et al. Bonsai: an event-based framework for processing and controlling data streams. Frontiers in neuroinformatics 9, 7, doi:10.3389/fninf.2015.00007 (2015).

40. Berman, G. J., Choi, D. M., Bialek, W. & Shaevitz, J. W. Mapping the stereotyped behaviour of freely moving fruit flies. Journal of the Royal Society, Interface 11, doi:10.1098/rsif.2014.0672 (2014).

41. Tao, L., Ozarkar, S., Beck, J. M. & Bhandawat, V. Statistical structure of locomotion and its modulation by odors. eLife 8, doi:10.7554/eLife.41235 (2019).

42. Henderson, J. M. et al. Behavioural effects of parafascicular thalamic lesions in an animal model of parkinsonism. Behavioural brain research 162, 222–232, doi:10.1016/j.bbr.2005.03.017 (2005).

43. Franklin, G. P. K. B. J. The Mouse Brain in Stereotaxic Coordinates. 4th edn, (Academic Press, 2012).

